# A connection between two ancient and essential cellular processes, iron-sulfur protein biogenesis and fatty acid synthesis, in *Escherichia coli*

**DOI:** 10.1101/2025.06.16.659903

**Authors:** Soufyan Fakroun, Guillaume Bouvier, Marouane Libiad, Emmanuel Sechet, Emmanuelle Bouveret, Frédéric Barras, Sarah Dubrac

**Affiliations:** Institut Pasteur, Université Paris-Cité, CNRS UMR 6047, Department of Microbiology, Stress Adaptation and Metabolism in enterobacteria Unit, Paris, France; Institut Pasteur, Université Paris-Cité, CNRS UMR 3528, Department of Structural Biology and Chemistry, Structural Bioinformatics Unit, Paris, France

**Keywords:** Iron Sulfur Cluster Biogenesis, Fatty Acid Biosynthesis, Acyl Carrier Protein, *E*. *coli*

## Abstract

Iron-sulfur [Fe-S] clusters are ubiquitous cofactors of a wide array of structural and functional diverse proteins. Acyl Carrier Protein (ACP) is the universal factor required for fatty acid (FA) synthesis. In this study in *E. coli*, we demonstrated that [Fe-S] and FA biosynthesis pathways are coordinated processes, driven by a physical interaction between ACP and the ISC [Fe-S] biogenesis machinery. Using bacterial two-hybrid assays, co-purification and biochemical analyses, we demonstrated a molecular interaction between ACP and IscS, the ISC machinery cysteine desulfurase that provides sulfur for [Fe-S] cluster formation. Structural modeling and directed mutagenesis pinpointed the ACP-binding site in a region of IscS shared for interactions with other components of the ISC [Fe-S] biogenesis system. At the cellular level, ACP depletion was found to disrupt ISC-dependent [Fe-S] cluster biogenesis, diminishing the activity of key [Fe-S]-dependent regulators (IscR, FNR, NsrR) and enzymes (aconitase, biotin synthase). Our findings underscore a functional link between [Fe-S] cluster biogenesis and fatty acid metabolism with far-reaching unexplored intricacies of metabolic coordination and cellular homeostasis. Comparison with eucaryotic systems highlight a strong evolutive driving force towards a link between [Fe-S] cluster and fatty acid biosynthesis in all living systems.

**Importance:** Cellular functions rely on interconnected metabolic pathways, yet many regulatory links remain unexplored. Iron-sulfur [Fe-S] clusters are co-factors of proteins driving fundamental cellular processes, from respiration to gene regulation. Our study uncovers a direct connection between [Fe-S] cluster biogenesis and fatty acid biosynthesis. We demonstrate the molecular connection between these two essential cellular processes to lie within the interaction between the acyl carrier protein (ACP), a shuttle of fatty acid biosynthetic intermediates and IscS, the source of sulfur for [Fe-S] cluster assembly. Intriguingly, similar interactions between ACP and [Fe-S] building cysteine desulfurase have been observed in yeast and human models, yet resting on different molecular determinants. This points out the existence of a strong evolutive driving force towards establishing a link between [Fe-S] cluster and fatty acid biosynthesis in all living systems with far-reaching implications for metabolic coordination and cellular homeostasis.

## Introduction

[Fe-S] clusters are metallic protein cofactors essential in nearly all living organisms. [Fe-S] proteins are involved in a wide range of cellular processes, such as respiration, metabolism, gene expression, RNA modification, lipid degradation, cell envelope biogenesis, vitamin and cofactor biosynthesis (1). They facilitate a wide array of biochemical processes from electron transfer to redox and non-redox chemical catalysis. *Escherichia coli* is predicted to synthesize more than 180 [Fe-S] proteins, with great diversity both in terms of structure and function. Hence, biogenesis of [Fe-S] cluster-containing proteins is essential for cell survival. This process is carried out by multi-protein machineries and occurs in two steps: the assembly of the [Fe-S] cluster and its delivery to apoprotein targets (2, 3). Five [Fe-S] biogenesis machineries were described: ISC in bacteria, archaea and mitochondria, SUF in bacteria, archaea, chloroplasts and organelles, MIS and SMS in both bacteria and archaea, and NIF in bacteria (4–6). The ISC machineries found in bacteria and mitochondria share significant similarities in functioning and composition. They include a cysteine desulfurase (IscS in bacteria and NFS1 in eukaryotes), a scaffold (IscU in bacteria and Isu in eukaryotes), a ferredoxin (Fdx in bacteria and Yah1 in eukaryotes), a frataxin (CyaY in bacteria and Ftn in eukaryotes), a co-/chaperone duo (HscBA in bacteria and Hsp70/40 in eukaryotes). Briefly, the scaffold captures sulfur, from cysteine desulfurase, and iron, from an unidentified source, assembles the [Fe-S] cluster, which is subsequently transferred to apo targets *via* [Fe-S] cluster carriers (7, 8).

In mitochondria, ACP (Acyl Carrier Protein) interacts with the cysteine desulfurase NFS1 *via* ISD11, a member of the LYRM protein family characterized by a leucine-tyrosine-arginine (LYR) motif (9–11). Experimental evidences suggested that ACP enhances the ISC-dependent synthesis of [Fe-S] clusters (12). Interestingly, bacterial ACP was also reported to interact with the IscS cysteine desulfurase, despite the absence of ISD11 in bacteria. Indeed, ACP was found to co-purify with IscS (13) and later on, IscS was identified as a potential partner of ACP in a TAP-tag screen (14).

ACP provides acyl chains for the synthesis of phospholipids, lipopolysaccharides, biotin and lipoate. The latter is an essential cofactor for enzymes of the TCA cycle such as the pyruvate dehydrogenase (PDH) that is crucial for the energy homeostasis in both bacteria and eukaryotes (15, 16). To be active, ACP undergoes a post-translational maturation during which a phosphopantetheine group (4’-PP), derived from coenzyme A, is added to the side chain of a serine residue in position 36 of ACP. This modified form is known as holo-ACP, which represents the active form of the carrier, capable of binding a malonyl chain (from malonyl-CoA) through *trans*-esterification to the thiol group at the 4’-PP terminus (Fig. S1). This acylated-ACP then interacts with the enzymes of the FASII pathway (Fab enzymes) that perform acyl chain elongation. Structurally, ACP is a small three-α-helix bundle protein. The serine 36 is located in the α_2_ helix, which mediates interactions with the Fab enzymes set (17). Beyond its interaction with the cysteine desulfurase IscS, ACP has also been found to interact with SpoT, a (p)ppGpp hydrolase, and MukB, a protein involved in chromosome partitioning during cell division (14, 18).

In the present study, we investigated the molecular basis and the physiological significance of the interaction between IscS and ACP. By using a biomolecular interaction modeling (Boltz) as well as bacterial two-hybrid assay, we identified key residues involved in the IscS/ACP interaction. Moreover, we demonstrated that ACP acts as a positive effector in the biogenesis of [Fe-S] clusters, specifically through the ISC system. Last we identified a series of [Fe-S] cluster bound protein targets, whose maturation efficiency was influenced by the amount of ACP in the cell. We propose that ACP availability (and/or its acylation state) helps to coordinate [Fe-S] cluster and fatty acid homeostasis, thereby adapting the efficiency of key cellular pathways to meet with cellular needs.

## Results

### A model-based prediction of the IscS-ACP interaction

Occurrence of an interaction between ACP and IscS was suggested by protein-protein interaction methods (13, 14). Therefore we wished to investigate the validity of this interaction by modeling the ACP-IscS complex using Boltz-1 (19). Resulting model showed that a dimer of IscS interacts with two ACP monomers (Fig. 1A). The phosphopantetheine group of holo-ACP is predicted to fit into a hydrophobic tunnel formed at the interface between the two IscS monomers (Fig. 1B and C). It is noticeable that the PLP (pyridoxal 5′-phosphate) cofactor of IscS also locates in this tunnel while the catalytic residue Cys_328_ is oriented toward the 4’-PP arm (Fig. 1B). Moreover, a positively charged region of IscS is predicted to interact with a negatively charged region of ACP, involving the pairs Arg_112_ and Arg_116_ of IscS with the Asp_35_ and Asp_38_ of ACP respectively (Fig. 1D). This analysis gave further support to the notion that *E. coli* IscS and ACP interact.

**Figure 1:**
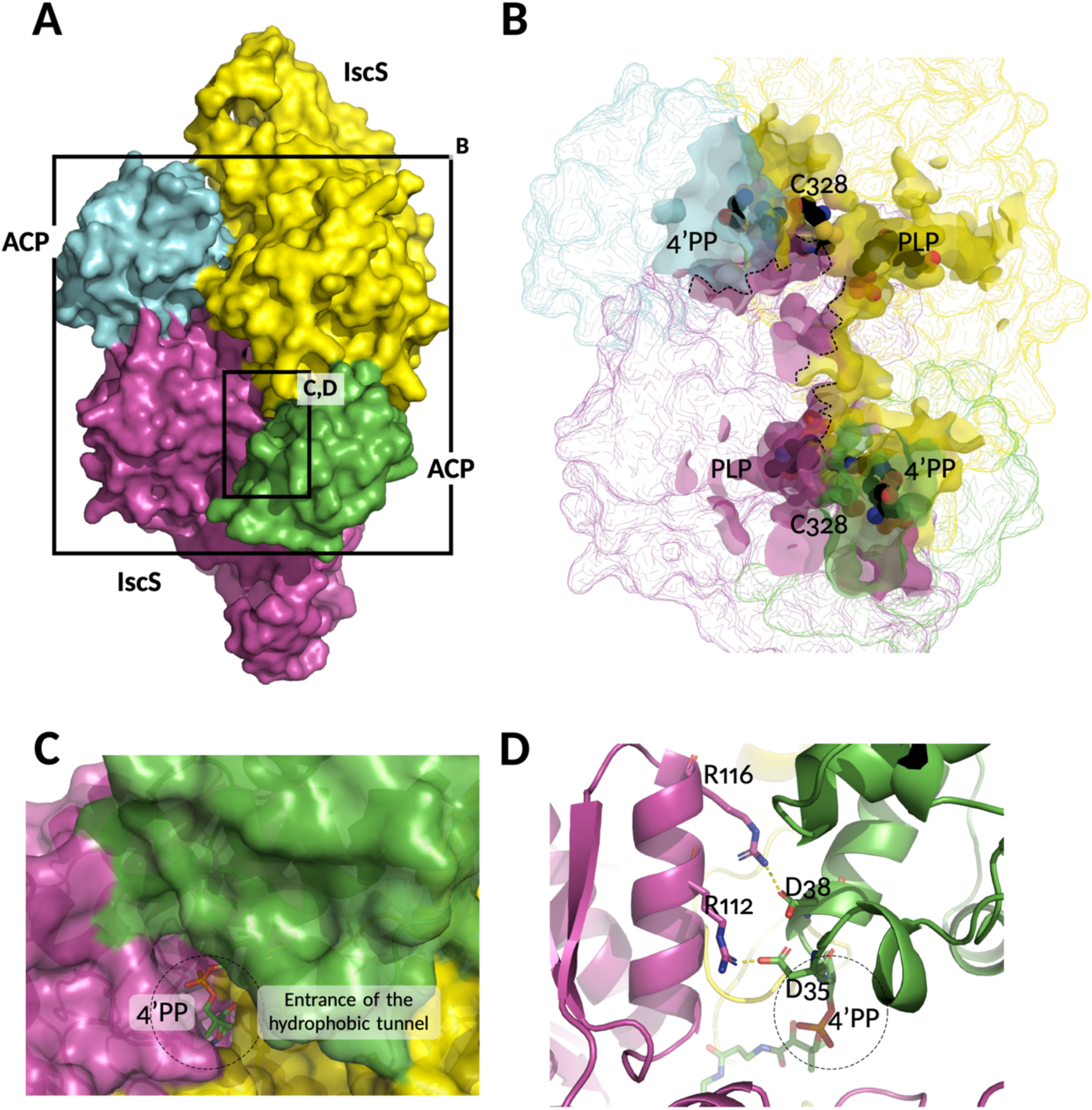
Modeling of the ACP-IscS complex. **A)** Overall structure of the ACP_2_-IscS_2_ complex showing surface interaction between an IscS dimer (magenta and yellow) and two ACP monomers (blue and green). Frameworks delineate the region of the structure represented in B, C and D as indicated. **B)** Inner view of the hydrophobic tunnel formed at the interface (dotted line) of the two IscS monomers. The 4’-PP arm is penetrating the tunnel, where the PLP is also present. The catalytic cysteine residue of IscS is also indicated. The tunnel is depicted in surface representation while the complex is shown as contour, keeping the same orientation as in A. **C)** Zoom in the ACP-IscS interaction zone shown as surface with the 4’-PP arm fitting into the hydrophobic tunnel at the interface between the two IscS monomers. **D)** Salt-bridge interactions detected at the interface between the ACP (green) and the IscS protein (magenta). Key amino acids involved in the interaction are shown as sticks, highlighting ACP residues Asp_35_ and Asp_38_, and IscS residues Arg_112_ and Arg_116_. The ACP-bound 4’-PP arm is also indicated and the entrance of the hydrophobic tunnel is outlined with a dotted-ring.

### Molecular characterization of the IscS-ACP interaction

In order to assess the model-based prediction above, the interaction between ACP and IscS was investigated using a series of different molecular approaches. Firstly, we used the Bacterial Adenylate Cyclase Two-Hybrid (BACTH) system (20). ACP was fused with the T25 subunit of *Bordetella pertussis* adenylate cyclase, while IscS was fused with the T18 subunit. Co-transformation of *E. coli* BTH101 cells with these constructs (T25-ACP and T18-IscS) resulted in high level of β-galactosidase activity (Fig. 2A and B), indicating an interaction between ACP and IscS. Two types of specificity controls were performed. Firstly, we took advantage of the capacity of human ISD11 to bind bacterial ACP (21) and reasoned that overexpression of ISD11 should compete with IscS for binding to ACP. We co-expressed human ISD11 and indeed observed a loss of interaction between ACP and IscS (Fig. 2A). Secondly, we tested whether SufS, another *E. coli* cysteine desulfurase, was able to bind ACP. No interaction was observed between T25-ACP and T18-SufS, supporting the notion that the interaction between ACP and IscS is specific (Fig. 2A).

**Figure 2:**
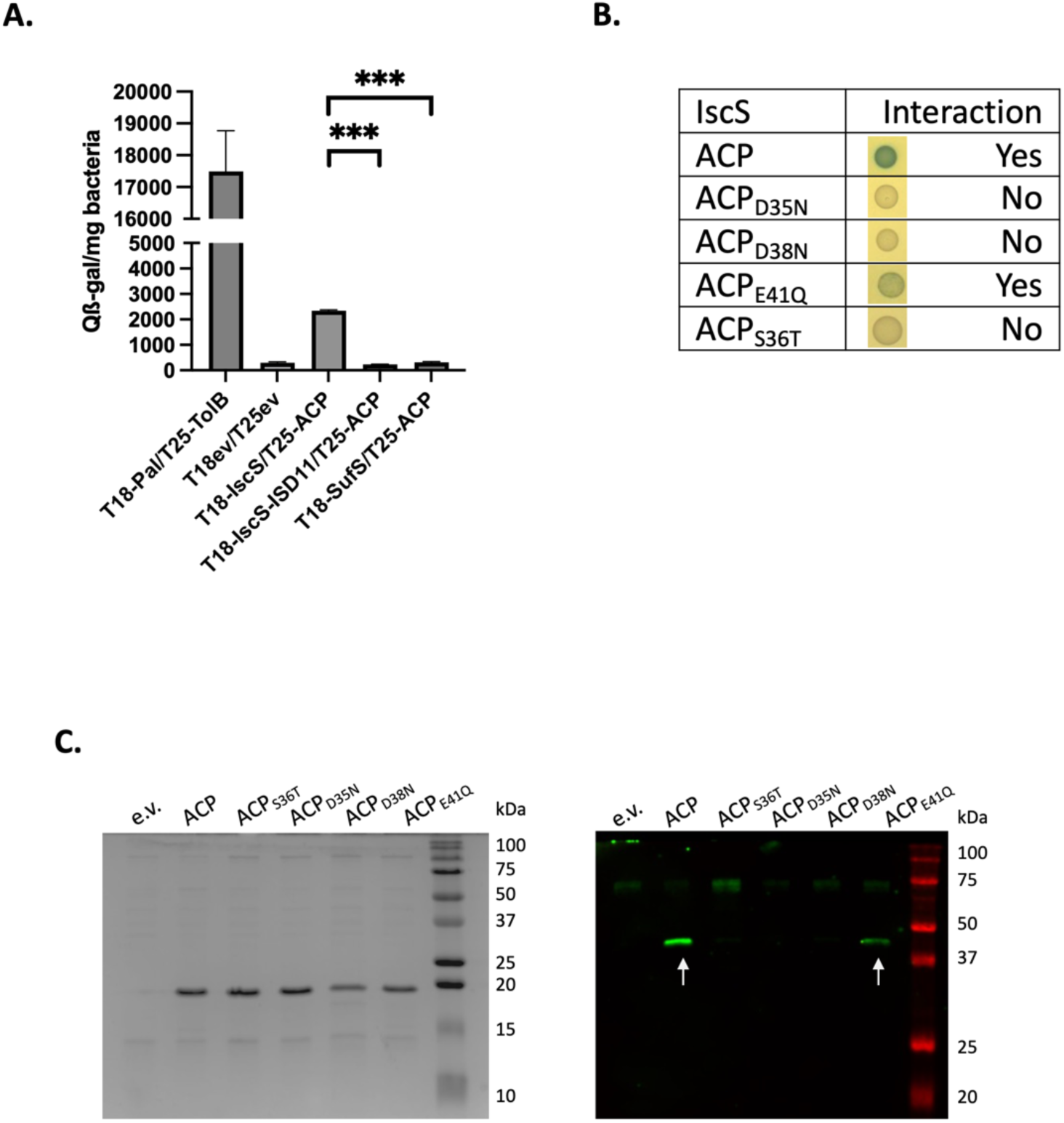
The IscS cysteine desulfurase forms a complex with ACP. **A)** The ACP protein was tested for pairwise interactions with the IscS and SufS cysteine desulfurases using BACTH assay. Translational fusions of tested proteins with T25 or T18 adenylate cyclase domains were co-transformed into BTH101 strain and ß-galactosidase activity assay for indicated pairs were performed. For testing ISD11 interferences, the human *isd11* was co-expressed with the T18-IscS fusion. A positive control of interaction using the Tol and Pal proteins was included as well as a negative control with a strain containing the pT18 and pT25 empty vectors. Data are the mean of data obtained from three biological replicates and error bars are standard errors of the mean. ***, P<0.001, one-way ANOVA with post hoc multiple comparisons performed via the Dunnett test. **B)** Role of negative residues of ACP (Asp_35_, Asp_38_, Asp_41_), as well as the phosphopantetheine binding site of ACP (Ser_36_) in ACP-IscS interactions. Fusions of ACP variant alleles (D_35_N, D_38_N, E_41_Q, S_36_T) with T25 were obtained by PCR site-directed mutagenesis of the parent pT25-ACP plasmid. BTH101 cells co-transformed with pT18-IscS and pT25-ACP and ACP variants were dropped (3 µL) on LB plates containing X-gal and IPTG and incubated for 24h at 30°C. **C)** Each pairwise interaction was confirmed using co-purification assay. CBP-ACP (or ACP variants as indicated) were produced from plasmids derived from the pBAD-CBP vector. Plasmids were transformed in the MG1655 strain. Resulting strains were grown in LB until OD_600nm_ 0.5 and exposed to 0.01% arabinose for 60 min at 37°C. After protein extraction and CBP protein fusions purification (see materials and methods), samples were analyzed on a 12% SDS-PAGE (left panel) showing CBP fusion proteins, and by Western-blot with anti-IscS antibodies (right panel). White arrows indicate IscS detection.

Next, using site directed mutagenesis, we generated variants of ACP that were modified on the residues predicted to be involved in the ACP-IscS interaction. The T25-ACP_D35N_ and T25-ACP_D38N_ were generated and tested for interaction with IscS using BACTH assay. We also included the T25-ACP_E41Q_ variant as a control to assess the effect of modifying another negative residue in the same region. Both T25-ACP_D35N_ and T25-ACP_D38N_ failed to bind to IscS, while a residual level of β-galactosidase activity was observed for the T25-ACP_E41Q_/T18-IscS pair (Fig. 2B). Additionally, the T25-ACP_S36T_ variant, in which the critical serine (S) residue at position 36 was changed to threonine (T), showed no interaction with IscS (Fig. 2B). This observation suggested that the serine in position 36 of ACP contributes to the interaction with IscS, presumably by allowing the 4’-PP arm to interact with IscS, as predicted by the model above.

Last, we used a co-purification based approach. A Calmodulin Binding Protein (CBP)-tagged ACP recombinant protein was produced from a plasmid. Both wild-type CBP-ACP and CBP-ACP variants were purified from cells (Fig. 2C, left panel). Endogenous IscS was co-purified with wild-type CBP-ACP and with the CBP-ACP_E41Q_ variant, but with none of the CBP fusions with ACP_D35N_, ACP_D38N_, or ACP_S36T_ variants (Fig. 2C, right panel). Consistent with the BACTH data, these results strongly supported the notion that ACP D_35_, D_38_, and S_36_ residues are critical for its interaction with IscS.

In parallel, we purified 6His-tagged versions of ACP (wild-type, D_35_N, and S_36_T) along with 6His-IscS using nickel affinity chromatography (Fig. 3A). Mixture of IscS with these different ACP were analyzed by native PAGE and Coomassie Blue staining. The migration pattern of IscS alone revealed several bands, likely corresponding to different oligomeric forms of IscS (Fig. 3B, first lane). Incubation of IscS with wild-type ACP resulted in a shift in the migration pattern, indicating interaction between IscS and ACP (Fig. 3B, white arrows). In contrast, no such shift was observed when IscS was incubated with the ACP S_36_T or D_35_N variants, further confirming the loss of interaction between these mutants and IscS (Fig. 3B). Collectively, these experiments demonstrated a direct and specific interaction between ACP and IscS.

**Figure 3:**
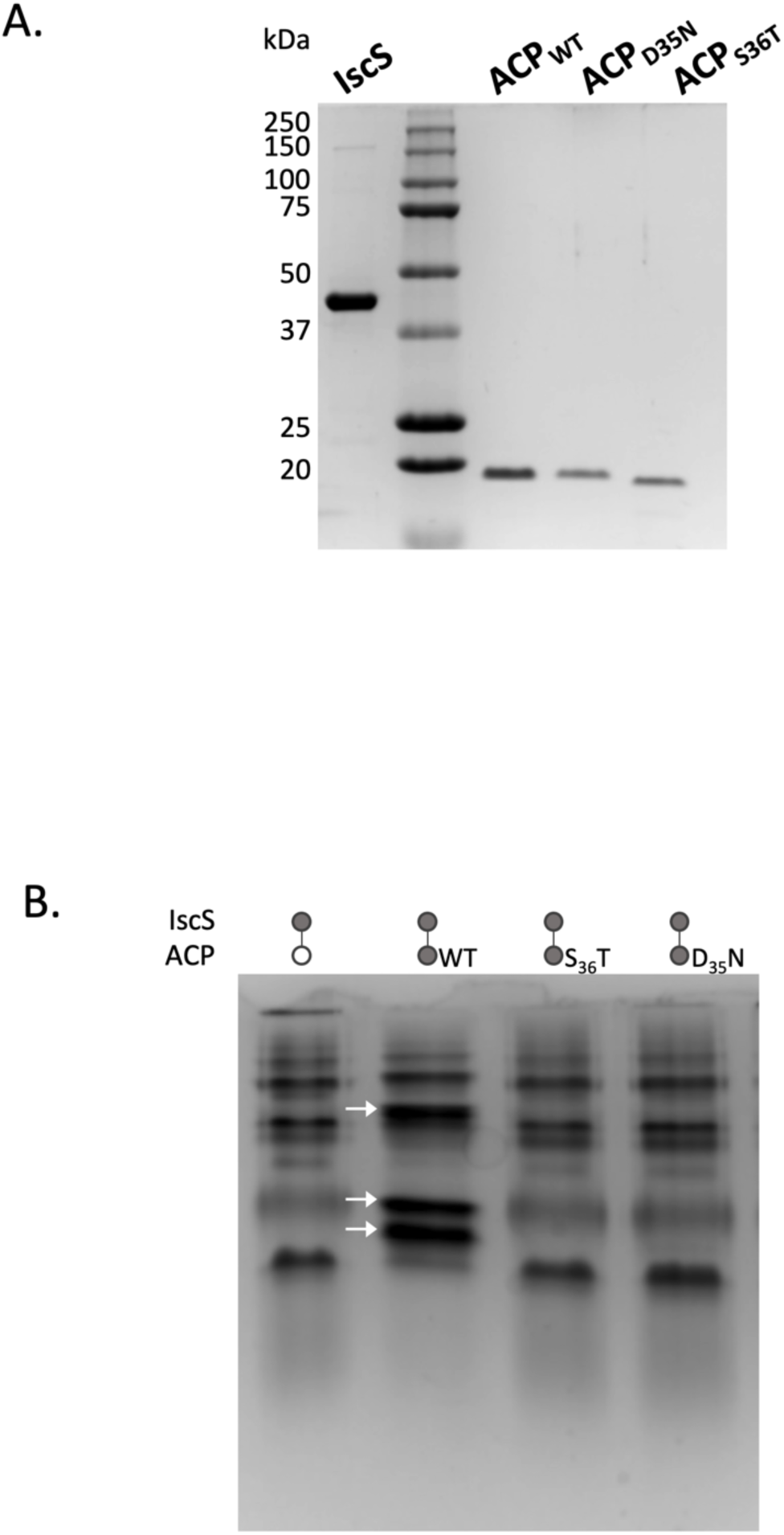
Direct *in vitro* interaction between ACP and IscS. **A)** SDS-PAGE of the purified 6His-IscS, 6His-ACP, and the ACP variants 6His-ACP_D35N_, and 6His-ACP _S36T_. **B)** Native PAGE showing 6His-IscS alone (15 µM) or mixed with 6His-ACP (30 µM for either WT, D_35_N, or S_36_T variants) as indicated. Prior to electrophoresis, protein mixtures were incubated in purification buffer for 20 minutes at 37°C (see Materials and Methods section). Loaded protein solutions are indicated above the gel picture. White arrows indicate shifted IscS bands upon incubation with ACP_WT_.

### Acylated ACP enhances the cysteine desulfurase activity of IscS

We next investigated whether ACP had an impact on the cysteine desulfurase activity of IscS. The purified ACP preparation was expected to comprise a mixture of apo-ACP, holo-ACP (phosphopantetheine-bound ACP), and a minor fraction of acylated ACP (22). Therefore, by using purified acyl-ACP synthetase (Aas), we acylated ACP with palmitic acid (C16:0). Upon SDS-PAGE analysis, the acylated ACP (ACP-C16) displayed a faster migration compared to the unmodified ACP, indicating successful acylation (Fig. 4A) (23). Only a portion of the purified ACP could be acylated, likely due to the proportion of holo-ACP available in the ACP purification. Desulfurase activity of IscS is increased 3-fold upon incubation with ACP-C16 whereas non-acylated form of ACP failed to enhance IscS activity (Fig.4B). These results showed that acylated-ACP enhances IscS cysteine desulfurase activity.

**Figure 4:**
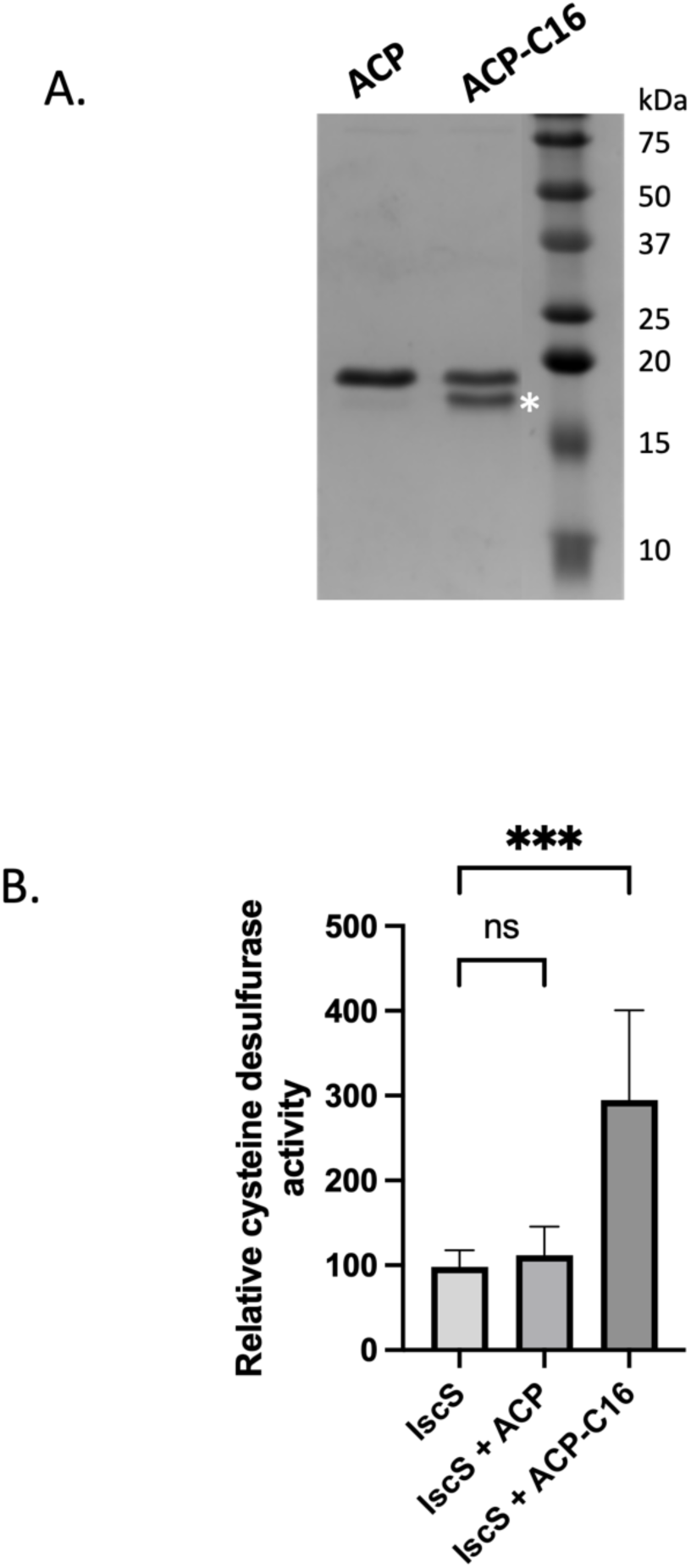
Acylated ACP enhances the cysteine desulfurase activity of IscS. **A)** Purified 6His-ACP was acylated using palmitate (C16:0) and pure Aas acyl-ACP synthase to catalyze the acylation. SDS-PAGE shows the purified 6His-ACP and acylated 6His-ACP, with the acylated form indicated by an asterisk. **B)** Cysteine desulfurase (CD) activity was measured using the methylene blue assay. CD activity was assessed for IscS alone and for IscS pre-incubated with either purified 6His-ACP or acylated 6His-ACP as described in A). Results are presented as relative CD activity normalized to the activity of IscS alone. Values and error bars represent the means and standard errors of the mean of six independent measurements. Statistical significance was determined using one-way ANOVA with post hoc multiple comparisons performed via the Dunnett test; ns: not significant; ***, P<0.001.

### Decrease in ACP levels leads to [Fe-S] dependent phenotypes

The *acpP* gene, which encodes ACP, is essential, thereby precluding the use of null mutant to investigate the effect of ACP levels on [Fe-S] cluster biogenesis *in vivo*. Therefore, we used CRISPR-based RNA interference (CRISPRi) to down regulate ACP levels. The CRISPRi system used in this study consisted of two plasmids: 1) the psgRNA plasmid, which produces a guide RNA under the control of a constitutive promoter, and 2) the pdCas9 plasmid, wherein the expression of dCas9 is controlled by the TetR-regulated promoter, Ptet, and can be induced by anhydrotetracycline (AnTet) (24). We designed a guide RNA (gRNA-*acp*) complementary to a portion of the *5’UTR* of *acpP* to generate the psgRNA-*acp* plasmid (Fig. S2A). To clarify, “e.v.” refers to strains containing the pdCas9 and psgRNA plasmids, while “CRISPRi-acp” indicates strains that carry the pdCas9 and psgRNA-*acp* plasmids. Induction of dCas9 expression with 500 ng/mL of AnTet prevented bacterial growth of the CRISPRi-acp strain (Fig. S2B). Therefore, we reduced the concentration of the dCas9 inducing AnTet down to 0.1 ng/mL, in order to allow bacterial growth. This yielded to a 5-fold decrease in *acpP* expression in the CRISPRi-acp strain compared to the e.v. control (Fig. S2C).

Next, we investigated the effect of reduced ACP levels on [Fe-S] cluster biogenesis in a *ΔsufBCD* strain, wherein [Fe-S] cluster formation is entirely dependent on the ISC pathway. We observed that the growth rate of the *ΔsufBCD/*e.v. strain was 66.55 ± 1.58 minutes, while the *ΔsufBCD/*CRISPRi-acp strain exhibited a doubling time of 85 ± 4.51 minutes. Conversely, lowering ACP levels had no noticeable effect in a wild-type background (MG1655, WT) as doubling times were 61 ± 1 and 67 ± 4 minutes for WT/e.v. and WT/CRISPRi-acp respectively (Fig. 5A). Next, we assessed the sensitivity of WT/e.v. and WT/CRISPRi-acp to aminoglycoside antibiotics, a phenotype that is controlled by the ISC system efficiency (25). The e.v. strain exhibited high sensitivity to gentamicin, with only 2% survival after 3h of treatment with 5 µg/mL of gentamicin. In contrast, the CRISPRi-acp strain showed enhanced tolerance, with over 8% survival, suggesting that reduced ACP levels impaired ISC activity, leading to enhanced aminoglycoside tolerance (Fig. 5B, right panel). As a control, we also tested a *ΔiscS* strain, which showed high level of tolerance (Fig. 5B, left panel). Overall, these results indicated that a reduction in ACP levels negatively affected ISC efficiency.

**Figure 5:**
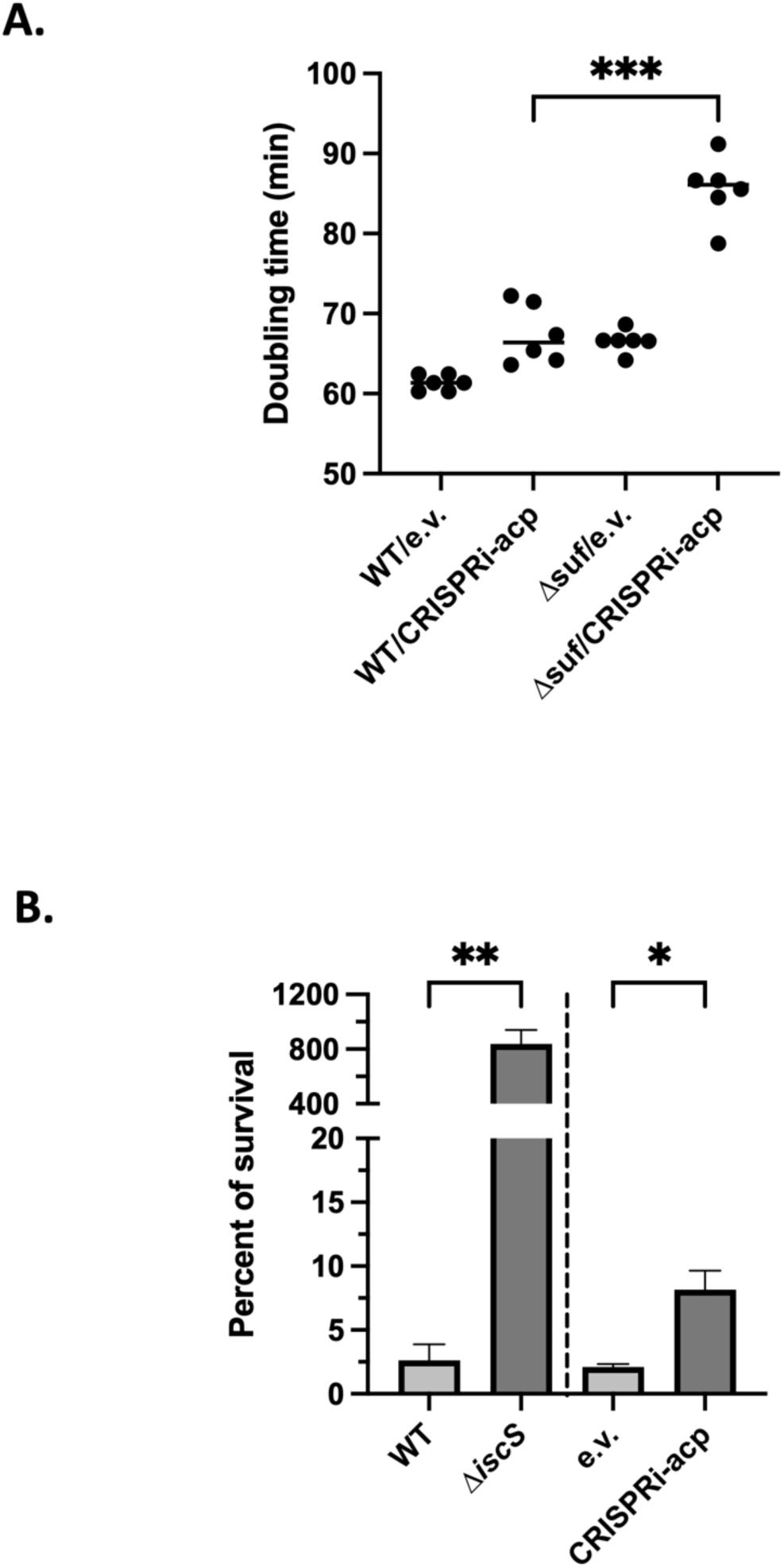
Reduced ACP levels affect the efficiency of the ISC machinery. **A)** Growth of WT/e.v., WT/CRISPRi-acp, a Δ*sufBCD*/e.v. and Δ*sufBCD*/CRISPRi-acp strains were done in LB medium with 0.1 ng/mL AnTet, and OD_600nm_ were measured using a Tecan microplate reader. WT was MG1655 strain. Doubling times were calculated during the exponential growth phase. Results of six biological replicates are presented with average growth rates and their respective variations. ***, P<0.001, unpaired Student’s t test. **B)** Strains were grown in LB for MG1655 (WT) and Δ*iscS* and LB-AnTet 0.1 ng/mL for the e.v. and CRISPRi-acp strains until OD_600nm_ around 1. Survival of strains was measured after 3 hours of treatment with gentamycin (5 µg/mL) by counting colony-forming units (CFU) and normalized to time at which the antibiotic was added to calculate the rate of survival. Data presented are the means of four biological replicates. **, P<0.01; *, P<0.05, unpaired Student’s t test.

### Transcriptional activity of [Fe-S]-dependent regulators is stimulated by ACP

To further assess the impact of reduced ACP levels on [Fe-S] cluster biogenesis *in vivo*, we tested if the activity of several well-characterized [Fe-S] cluster regulators was affected by reducing the ACP levels. Specifically the activity of IscR, FNR, and NsrR which are [Fe-S] bound transcriptional regulators involved in [Fe-S] homeostasis, anaerobic adaptation, and nitrosative stress responses, respectively, were assayed (26).

IscR is active under two forms: the holo-IscR (containing a [2Fe-2S] cluster), which represses its own expression among other targets, and the apo-IscR, which regulates another set of genes (27). We utilized a P*iscR*-*lacZ* chromosomal fusion to monitor the efficiency of IscR maturation. In strains grown in rich LB medium, the P*iscR*-*lac*Z fusion exhibited less than 20 units/mg of β-galactosidase supporting the notion that IscR was predominantly in its repressing holo-form (Fig. S3). Upon addition of 2,2’-dipyridyl (DIP), an iron chelator, expression increased in a dose-dependent manner, reflecting a shift from the holo-to the apo-form of IscR and the associated alleviation of repression (Fig. S3). The prediction was that lowering the level of ACP should lower the efficiency of ISC mediated maturation of IscR, which will eventually lead to the de-repression of the P*iscR*-*lacZ* fusion. The experiment was run in the presence of 62.5 µM DIP, a condition in which Fe-limitation is modest and likely allows the presence of both apo- and holo-forms of IscR. In this condition, depletion of ACP *via* CRISPRi (CRISPRi-acp) resulted in more than 2-fold increase in P*iscR*-*lacZ* expression (Fig. 6A, dark grey plots). Importantly, in a Δ*iscR* strain, P*iscR*-*lacZ* expression was significantly higher, reflecting the absence of the IscR repressor. It was also insensitive to changes in ACP levels, demonstrating that the ACP-mediated effect on *iscR* expression was IscR-dependent (Fig. 6A, light grey plots).

**Figure 6:**
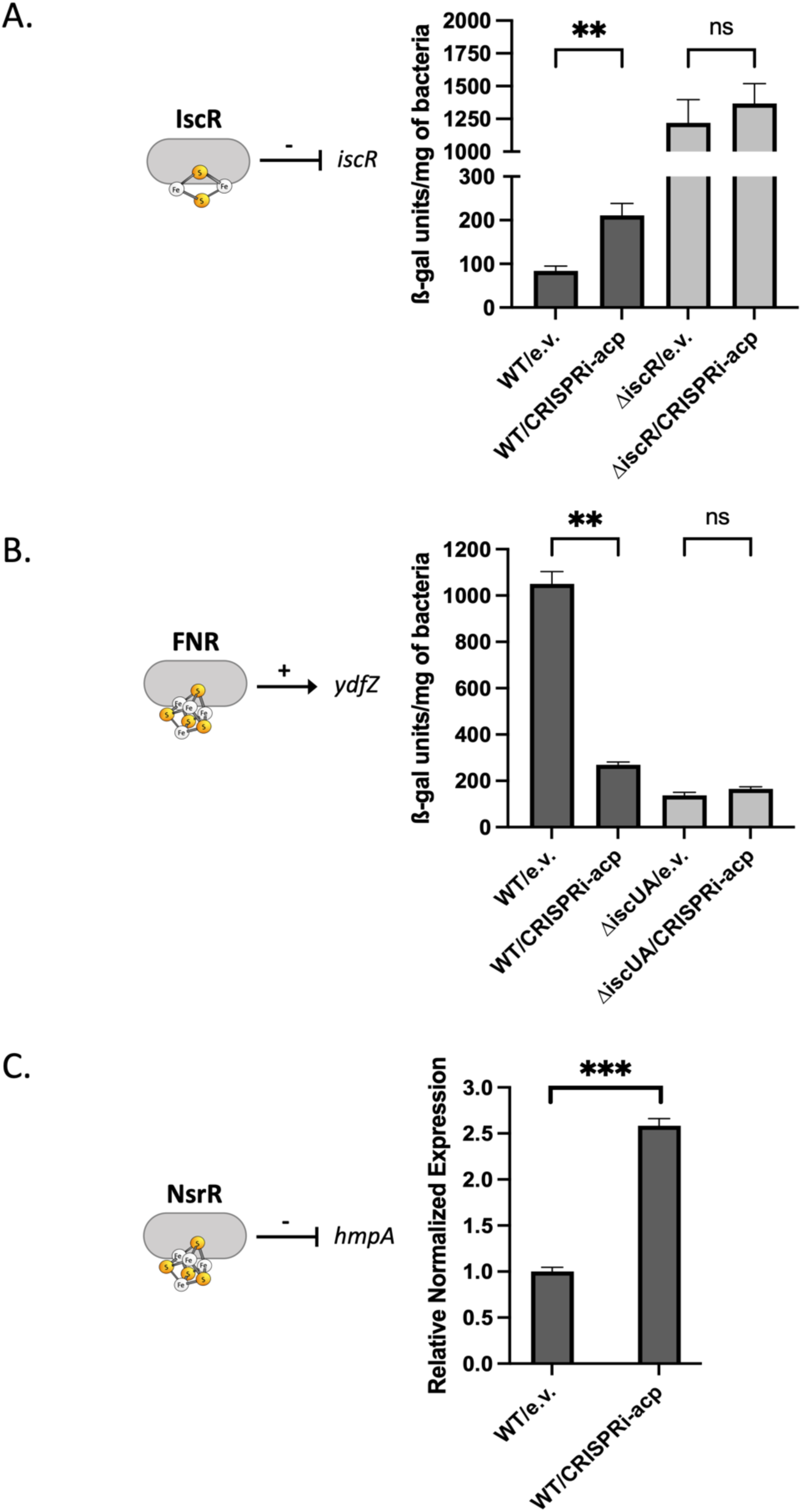
ACP levels interfere with the activity of [Fe-S] regulators. **A)** Activity of the *iscR* promoter was assessed by measuring the ß-gal activity (Miller units) of the P*iscR-lacZ* strains carrying pdCas9 and psgRNA (e.v.) or pdCas9 and psgRNA-*acp* (CRISPRi-acp) in either a WT (FBE005) background, or a Δ*iscR* (FBE008) background. Strains were grown in LB with 0.1 ng/mL of AnTet and 62 µM of dipyridyl until OD_600nm_ = 2 before β-gal activity was measured. **B)** Activity of the *ydfZ* promoter was assessed by measuring the β-gal activity (Miller units) of the P*ydfZ-lacZ* strains carrying the indicated plasmids in either a WT background (FBE204), or a Δ*iscUA* background (FBE596). Strains were grown in LB with 0.1 ng/mL of AnTet until OD_600nm_ = 2 before β-gal activity was measured. **C)** Expression of *hmpA* in a WT (MG1655) background carrying pdCas9 and psgRNA (e.v.) or pdCas9 and psgRNA-*acp* plasmids (CRISPRi-acp). Strains were grown in LB with 0.1 ng/mL of AnTet until OD_600nm_ = 1 before RNA extraction. Gene expression was measured using quantitative Real Time PCR (qRT-PCR). Expression levels were normalized using 16S rRNA as internal standard and presented as the n-fold change of the CRISPRi-acp strain compared to the e.v. strain. Results are presented as the mean of 4 replicates and standard errors of the mean are indicated. Statistical significance was determined using one-way ANOVA with post hoc multiple comparisons performed via the Dunnett test; ***, P<0.005; **, P<0.01; ns, no significative difference.

FNR is active upon binding a [4Fe-4S] cluster and is inactivated by oxygen (28). We measured P*ydfZ*-*lacZ* activity as a proxy for FNR activity. The *ydfZ* promoter remains activated by FNR under aerobic conditions, albeit at a reduced level (29, 30). While the WT strain exhibited high levels of β-galactosidase activity, depletion of ACP led to a significant decrease in activity, reducing it by approximately 5-fold (Fig. 6B, dark grey plots). In contrast, in a *ΔiscUA* background, where FNR maturation is impaired, P*ydfZ-lacZ* activity was reduced compared to the WT, and varying ACP levels had no further effect on this strain (Fig. 6B, light grey plots). This result supported the notion that ACP depletion impacted ISC-dependent maturation of FNR.

NsrR is a [4Fe-4S] containing regulator that plays a crucial role in the bacterial response to nitrosative stress. Under its holo-form NsrR acts as a repressor of NO defense-related genes, such as *hmpA* (31). To investigate the impact of ACP on NsrR activity, we monitored *hmpA* expression by qRT-PCR. Upon reduced ACP amount (CRISPRi-acp), we observed a more than 2-fold increase in *hmpA* expression (Fig. 6C), consistent with the notion that ACP enhanced NsrR repression activity.

SoxR binds a [2Fe-2S] cluster. It is activated under exposure to redox cycling drugs such as paraquat through redox state change of its [Fe-S] cluster (from +1 to +2). Importantly, under paraquat exposure, SoxR acquires its [Fe-S] cluster exclusively through the SUF system (32, 33). Using a P*soxS*-*lacZ* transcriptional fusion as a reporter for SoxR activity in presence of paraquat, we observed that the induction of *soxS* expression was not altered in strain with reduced ACP levels (Fig. S4A). This showed that ACP depletion did not affect [Fe-S] biogenesis carried out by the SUF system.

Together, these results showed that ACP enhanced maturation of [Fe-S] cluster-containing regulators, such as IscR, FNR, and NsrR and strengthened the conclusion that ACP had a positive effect on ISC dependent production of [Fe-S] clusters. Importantly it had no effect on SUF dependent [Fe-S] cluster production.

### Activity of [Fe-S]-dependent enzymes is stimulated by ACP

To investigate whether ACP positive effect on ISC activity extends to [Fe-S] dependent enzymes, we analyzed the activity of aconitase and biotin synthase. Aconitase activity, which results from the activity of two [4Fe-4S] enzymes AcnA and AcnB, exhibited a 40% reduction in activity in a Δ*iscS* background, a result consistent with prior studies (Fig. 7A, left panel) (34). CRISPRi-mediated down-expression of *acpP* reduced aconitase activity by 30% compared to the empty vector control (Fig. 7A, right panel), highlighting a positive role of ACP in supporting aconitase activity.

**Figure 7:**
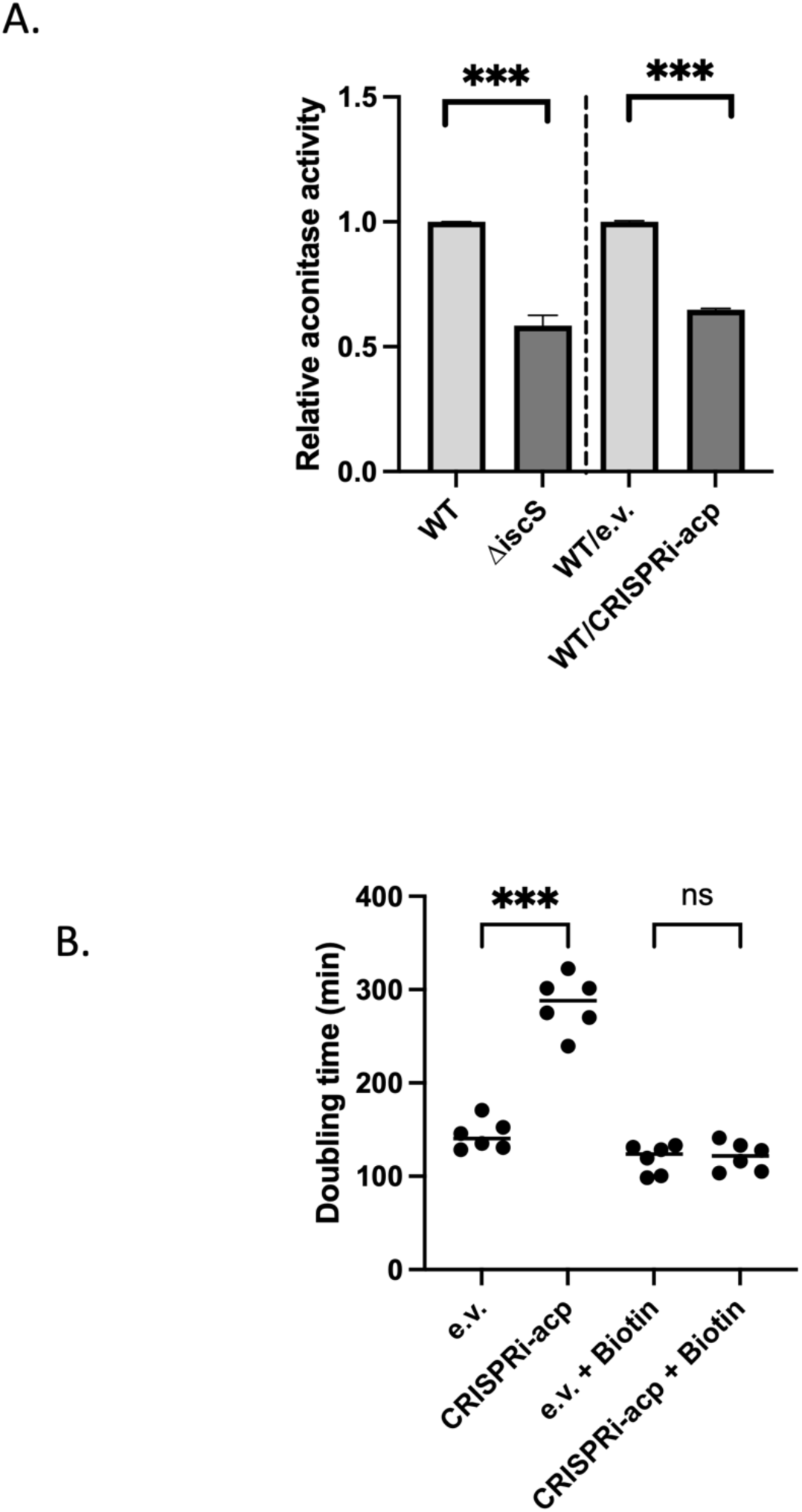
ACP affects activity of aconitase and biotin synthase [Fe-S] enzymes. **A)** Wild type strain (MG1655) and Δ*iscS* derivative were grown in LB, and wild type carrying pdCas9 and psgRNA (e.v.) or psgRNA-*acp* (CRISPRi-acp) were grown in LB with 0.1 ng/mL of AnTet, until OD_600nm_=2. Aconitase activity was measured on crude extracts immediately after sonication (see Materials and Methods). Results are presented as relative aconitase activity compared to either the wild type (left panel) or the wild type carrying pdCas9 and psgRNA (right panel). Values represent the mean of at least 4 biological replicates and standard errors of the mean are indicated. Statistical significance was determined using one-way ANOVA with post hoc multiple comparisons performed via the Dunnett test; ***, P<0.005. **B)** Bacterial growth rate was used as a read out of biotin synthase activity. Mutant strain with a deletion in *bioD* carrying pdCas9 and psgRNA (e.v.) or psgRNA-*acp* (CRISPRi-acp) were grown in M9 minimal medium supplemented with dethiobiotin (0.1 mM) as a substrate of BioB. Biotin (0.1 mM) was added in half of the cultures where indicated. Growth rates are presented as mean doubling time of six biological replicates for each strain and condition. Statistical significance was determined using one-way ANOVA with post hoc multiple comparisons performed via the Dunnett test; ***, P<0.005; ns, no significative difference.

BioB is a [2Fe-2S] enzyme essential for the final step of synthesis of biotin, an essential vitamin (Fig. S5 A and B). In a Δ*bioD* strain, where biotin synthesis depends on BioB-catalyzed conversion of dethiobiotin (DTB) to biotin, fitness in DTB-supplemented minimal medium served as an indirect measurement of BioB activity (35). Down-regulation of *acpP* in the Δ*bioD* background resulted in a doubling time of 288 minutes, a 2-fold increase compared to the empty vector strain (140 minutes) in minimal medium with DTB. This fitness defect was fully rescued by supplementing biotin, confirming that reduced growth was due to impaired BioB activity (Fig. 7B). Importantly, ACP knockdown did not affect the activity of a non-[Fe-S] enzyme, *i.e.* endogenous β-galactosidase (Fig. S4B). Together, these results demonstrated that ACP is a positive effector of [Fe-S]-dependent enzymes aconitase and biotin synthase, likely by enhancing their maturation.

### tRNA thiolation is not altered by varying ACP levels

IscS also acts as a sulfur donor for tRNA modification (36). Therefore, we investigated whether ACP also influences IscS-dependent tRNA thiolation catalyzed by MnmA, TtcA, MiaB and ThiI. Our results indicated no significant changes in tRNA thiolation levels under ACP depletion (Fig. S6). Whether the lack of an observable effect is due to insufficient impairment of IscS activity under ACP depletion or because the influence of ACP is specifically targeted toward the [Fe-S] biogenesis pathway remains to be established.

## Discussion

Our study revealed a connection between two essential and ancient cellular processes, fatty acid metabolism and [Fe-S] cluster biogenesis. We identified the molecular basis of this connection by uncovering a physical interaction between ACP, the acyl carrier protein that provides acyl chain to fatty acid biosynthetic pathway, and IscS, the cysteine desulfurase that provides sulfur for [Fe-S] biogenesis. We showed that ACP levels directly influence efficiency of cellular [Fe-S] bound proteins, whether they are transcriptional regulators, metabolic enzymes or respiratory chain components.

Using a combination of Boltz modeling, bacterial two-hybrid assays, co-purification experiments, and in vitro binding assays, we demonstrated that ACP interacts with the IscS cysteine desulfurase, enhancing its enzymatic activity. ACP is initially produced as an apo-form and undergoes two maturation steps. It is first modified by the covalent attachment of a 4’-phosphopantetheine group to its serine residue at position 36 (holo-ACP), and subsequently acylated with a malonyl moiety. While as-purified ACP, which is expected to comprise equal proportions of apo- and holo-forms and a small fraction of the acylated form (22), binds to IscS *in vitro*, only the acylated form of ACP exerts a positive effect on IscS activity. Both the maturation state of ACP and the specific acyl chain attached to it are expected to reflect the activity of the fatty acid biosynthesis machinery (37). Hence, it is tempting to propose that the IscS/ACP interaction allows the level of available fatty acids to fine-tune [Fe-S] biogenesis efficiency and eventually serves as a strategy to coordinate the levels of fatty acids and [Fe-S] clusters.

Computational modeling, mutagenesis, and biochemical analyses suggested that the interaction between ACP and IscS involves negatively charged residues on ACP (Asp_35_ and Asp_38_) and positively charged ones on IscS (Arg_112_ and Arg_116_). Moreover, computational model indicates that the ACP attached phosphopantetheine group may fit into a hydrophobic pocket at the interface of two IscS monomers, thereby enhancing the stability of the IscS₂-ACP₂ complex. Consistently, the ACP Ser_36_ residue, which is positioned in the vicinity of Asp_35_ and Asp_38_ residues and carries the 4’-phosphopantetheine group, is crucial for ACP binding to IscS. Interestingly the triad Ser_36,_ Asp_35_, Asp_38_ is located within the same α2 helix that contains the binding site for fatty acid biosynthesis enzymes FabA, FabI, and FabG (38–40). Conversely, on the IscS side, the Arg_112_ and Arg_116_ residues locate within the region involved in the interaction with several IscS partners. In particular, the Arg_116_ is crucial for IscS binding to CyaY and IscX, two proposed effectors of ISC-dependent [Fe-S] biogenesis (41, 42). Taken together these observations indicates that ACP binds within a region of IscS that serves as a hub for complex and potentially mutually exclusive interactions between IscS and its multiple partners.

In mitochondria, ACP and NFS1 interact via an intermediate protein, ISD11, yielding to a stable ACP-ISD11-NFS1 complex (43). In the yeast model, ACP was also shown to potentiate [Fe-S]-dependent aconitase activity (12). By using the Foldseek search method we failed to identify structures similar to ISD11 within the AlphaFold database, including bacterial proteins (44). Additionally, a BLAST search using the ISD11 sequence yielded no matches in the *E. coli* sequence database and only one hit in prokaryotic databases: a LYR motif-containing protein in *Kocuria palustris*. We therefore concluded that a LYR motif-containing motif mediating NFS1/ACP interaction is not to be found in *E. coli*. Furthermore, our *in vitro* binding assays indicate that ACP directly binds to bacterial IscS, pointing to a major difference between the bacterial and eukaryotic situations. Yet, it is noteworthy that 2 out of 4 residues of NFS1 involved in interaction with ISD11 are conserved in IscS while ACP residues involved in interaction with ISD11 in eukaryotes are conserved in *E. coli* ACP (Fig. S7, blue stars) (10). The evolutive advantage, if any, of the presence of such residues in the absence of ISD11 remains unclear. Conversely, although the key residues of *E. coli* ACP required for its interaction with IscS are conserved in eukaryotes, only one of the two arginine residues from IscS involved in ACP binding is conserved in NFS1 (Fig. S7, red stars). This difference may explain why NFS1 is unable to bind ACP directly.

ACP plays a central role in fatty acid biosynthesis, acting as a carrier for fatty acids during their elongation within the FASII cycle. The question arises of what is the teleonomic advantage of coupling fatty acid biosynthesis and [Fe-S] biogenesis. In fact, there are several proteins whose role and/or function depend upon both fatty acid and [Fe-S] biology and which could benefit from a coordinated regulation of [Fe-S] cluster and fatty acid syntheses. A first convergent node is BioB an enzyme catalyzing synthesis of biotin, an essential vitamin. BioB is a radical SAM enzyme hosting a [4Fe-4S] and a [2Fe-2S] clusters. It intervenes at a last step of a pathway fed by ACP-bound intermediates (Fig. S5A) (35, 45). Our results showed that decreased ACP levels decreased BioB activity, suggesting that the positive effect of acylated ACP on [Fe-S] biogenesis could help to coordinate both the levels of [Fe-S] clusters and fatty acid precursors for optimizing BioB activity. A second convergent node is LipA, an enzyme catalyzing synthesis of lipoate, an essential cofactor. Like BioB, LipA is a radical SAM enzyme hosting two [4Fe-4S] clusters, acting downstream a series of steps involving ACP-bound intermediates (45). Having LipA activity being optimized by a coordination between level of [Fe-S] and precursors is a reasonable hypothesis. A third convergent node arises with IspG and IspH, two [4Fe-4S] enzymes, required for the synthesis of isopentenyl phosphate, precursor of lipid II and LPS (46). In this case coordination between fatty acids and [Fe-S] cluster biogenesis could be a way to synchronize cell elongation as both feed into membrane and cell wall biogenesis. In this context, it is noteworthy that ACP has been shown to interact with MukB and to be essential for its ATPase activity. MukB is a core component of the Structural Maintenance of Chromosomes (SMC) complex, which plays a crucial role in chromosome segregation during cell division (18, 47). Hence, our study revealing a coordination between ACP and another central process such as [Fe-S] biogenesis points to the possibility of ACP acting as a cellular hub integrating homeostasis-regulating signals to coordinate cell cycle progression, division, and proliferation.

## Materials and Methods

### Media and growth conditions

Bacterial strains were grown in LB (Lysogeny Broth) rich medium at 37°C under agitation. Antibiotics were added when necessary: ampicillin (Amp, 100 µg/ml), kanamycin (Kan, 50 µg/ml), chloramphenicol (Cm, 10 µg/ml) and anhydrotetracycline (AnTet) for inducible expression from the Ptet promoter. For iron chelation, 2,2′-Dipyridyl (DIP, 62 µM) was added to the culture medium. Paraquat was used at 100 µM.

### Bacterial strains and plasmids

*Escherichia coli* K12 MG1655 strain, its derivatives and plasmids used in this study are listed in Tables 1 and 2, respectively. *E. coli* K12 XL1B strain was used for cloning procedures. Primers, with their sequences and descriptions are listed in Table 3.

**Table 1:** Strains used in this study.

| Lab ref. | Name | Genotype | Reference |
| --- | --- | --- | --- |
| FBE051 | MG1655 | Wild type K12 *E. coli* | F. Barras, lab collection |
| FBE052 | XL1B | *recA1 endA1 gyrA96 thi-1 hsdR17 supE44 relA1 lac* [F' *proAB lacIqZΔM15 Tn10* (Tet<sup>r</sup>)] | Stratagene |
| FBE054 | BTH101 | F<sup>-</sup>, *cya-99, araD139, galE15, galK16, rpsL1* (Str<sup>R</sup>), *hsdR2, mcrA1, mcrB1, relA1* | (20) |
| FBE001 | *PsoxS-lacZ* | MG1655 $\Delta lacZ$ *PsoxS::lacZ* | B. Py |
| FBE005 | *PiscR-lacZ* | MG1655 $\Delta lacZ$ *PiscR::lacZ* | B. Py |
| FBE204 | *PydfZ-lacZ* | MG1655 $\Delta lacZ$ *PydfZ::lacZ* (Kan<sup>R</sup>) | B. Py |
| FBE227 | $\Delta iscS$ | MG1655 $\Delta iscS::kan$ | This work |
| FBE228 | BL21(DE3) | *E. coli B dcm ompT hsdS(r<sub>B</sub><sup>-</sup>m<sub>B</sub><sup>-</sup>) gal* (DE3) | (62) |
| FBE008 | $\Delta iscR$ *PiscR-lacZ* | MG1655 $\Delta lacZ \Delta iscR$ *PiscR::lacZ* | B. Py |
| FBE596 | $\Delta iscUA$ *PydfZ-lacZ* | MG1655 $\Delta lacZ \Delta iscUA$ *PydfZ::lacZ* (Kan<sup>R</sup>) | F. Barras, lab collection |

**Table 2:** Plasmids used in this study.

| Lab ref. | Name | Description | Reference |
| --- | --- | --- | --- |
| **CRISPR interference** |  |  |  |
| pEB1976 | pdCas9 | Cm<sup>R</sup>, p15A ori, Ptet-dCas9 | (24) |
| pEB1977 | psgRNA | Amp<sup>R</sup>, ColE1 ori, PJ23119 promoter | (24) |
| pSF4 | psgRNA-ACP | Generated by PCR with ebm1822-ebp8 | This study |
| **Protein purification** |  |  |  |
| pEB1188 | pET-6His-TEV | Amp<sup>R</sup>, ColE1 ori, T7 promoter | (57) |
| pSF19 | pET-6His-TEV-*iscS* | Insertion of OSD56-57 in pET-6His-TEV (*EcoRI/XhoI*) | This study |
| pSF51 | pET-6His-TEV-*acpP* | Insertion of OSD146-147 in pET-6His-TEV (*EcoRI/XhoI*) | This study |
| pSM61 | pET-6His-TEV-*acpP*<sub>D35N</sub> | Directed mutagenesis on pSF51 using OSD266-267 | This study |
| pEB1161 | pET-6His-TEV-*acpP*<sub>S36T</sub> | 6His-TEV-*acpP*<sub>S36T</sub> under the control of the T7 promoter | (23) |
| pEB0817 | pET28-*aas*-6His | *aas*-6His under the control of the T7 promoter | (58) |
| **BACTH** |  |  |  |
| pEB354 | pT25link | KmR, p15A ori, Plac | (51) |
| pEB355 | pT18link | AmpR, ColE1 ori, Plac | (51) |
| pEB375 | pT25-ACP | Translational fusion between T25 and ACP | (51) |
| pTFP43 | pT25-ACP<sub>D35N</sub> | Translational fusion between T25 and the ACP<sub>D35N</sub> allele | This study |
| pTSF18 | pT25-ACP<sub>S36T</sub> | Translational fusion between T25 and the ACP<sub>S36T</sub> allele | This study |
| pTFP22 | pT25-ACP<sub>D38N</sub> | Translational fusion between T25 and the ACP<sub>D38N</sub> allele | This study |
| pTFP42 | pT25-ACP<sub>E41Q</sub> | Translational fusion between T25 and the ACP<sub>E41Q</sub> allele | This study |
| pTSF33 | pT18-IscS-ISD11 | Translational fusion between T18 and *iscS* with ISD11 under the control of the Plac promoter | This study |
| pEB362 | pT25-TolB | Translational fusion between T25 and TolB | (51) |
| pEB541 | pT18-IscS | Translational fusion between T18 and *IscS* | This study |
| pEB594 | pT18-SufS | Translational fusion between T18 and *SufS* | This study |
| pEB356 | pT18-PAL | Translational fusion between T18 and PAL | (51) |
| **Protein co-purification** |  |  |  |
| pEB602 | pBAD24-CBPlink | AmpR, pBR322 ori, *Pbad* | (51) |
| pEB540 | pBAD24-CBP-ACP | Translational fusion between CBP and ACP | (14) |
| pEB797 | pBAD24-CBP-ACP<sub>S36T</sub> | Translational fusion between CBP and ACP<sub>S36T</sub> | (51) |
| pSD53 | pBAD24-CBP-ACP<sub>D35N</sub> | Translational fusion between CBP and ACP<sub>D35N</sub> | This work |
| pSD54 | pBAD24-CBP-ACP<sub>D38N</sub> | Translational fusion between CBP and ACP<sub>D38N</sub> | This work |
| pSD55 | pBAD24-CBP-ACP<sub>E41Q</sub> | Translational fusion between CBP and ACP<sub>E41Q</sub> | This work |

**Table 3:**
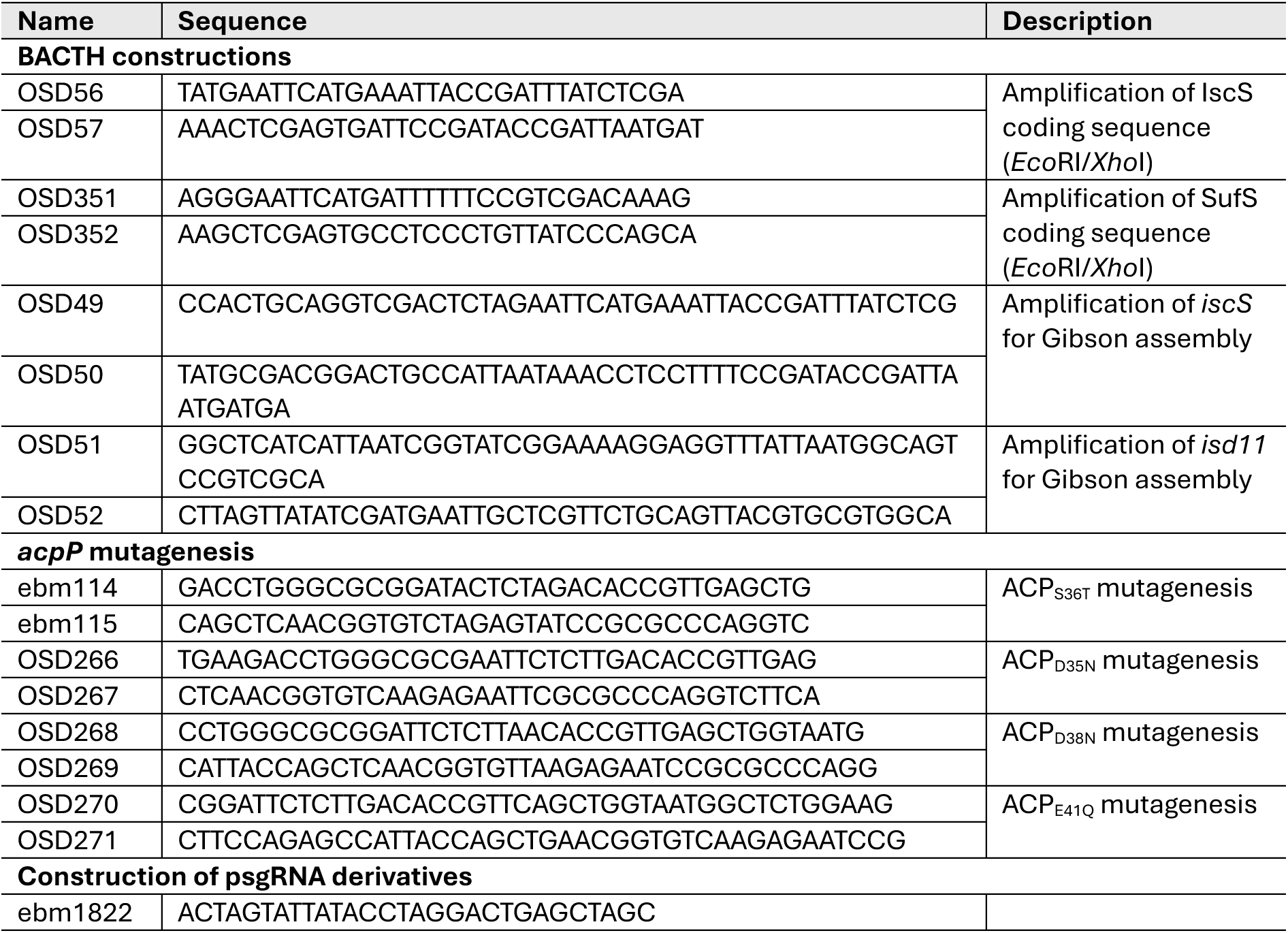
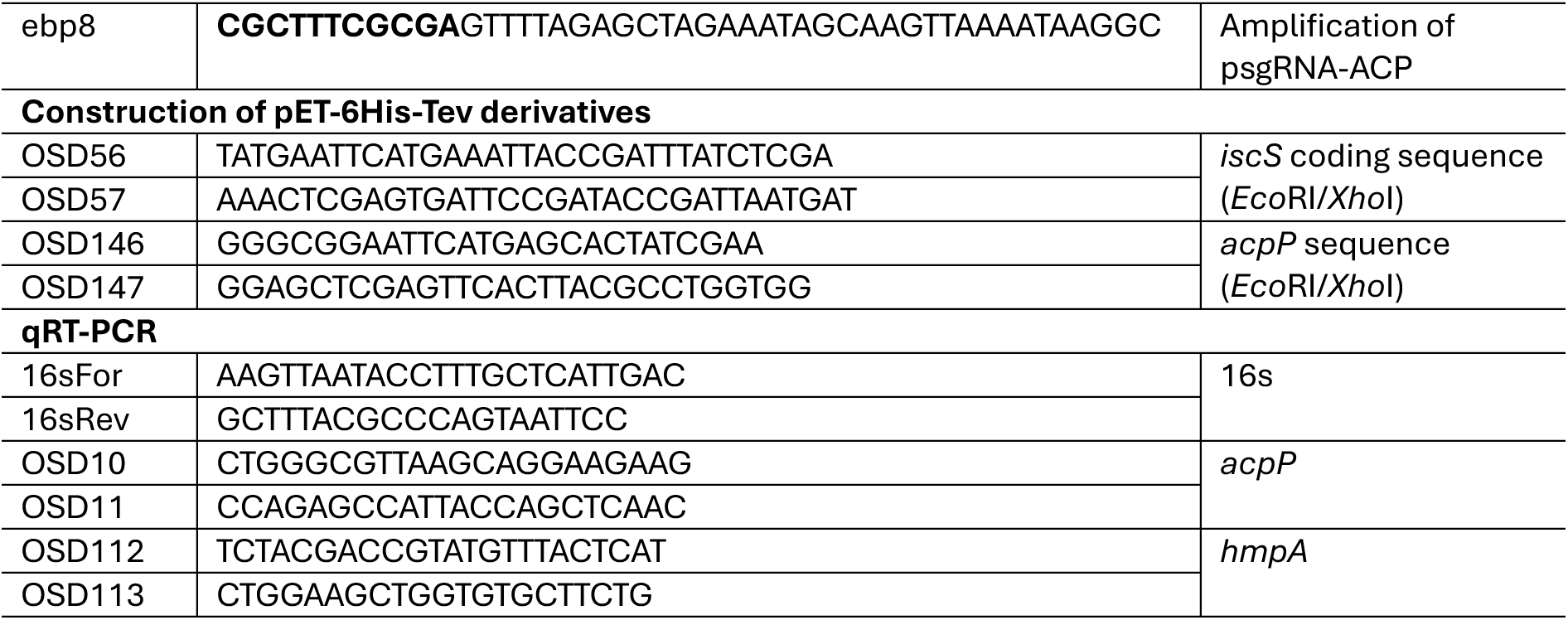
Primers used in this study.

Deletion mutations were introduced in MG1655 by P1 transduction using KEIO collection or lab strains as donor (48, 49). Transduced strains were verified by PCR using primers pair hybridizing upstream and downstream the deleted genes.

Standard procedures for preparation of DNA, amplification, digestion, ligation were used as previously described (50). Transformations were done by electroporation. Plasmids constructed for this study have been checked by sequencing (Eurofins).

Plasmid expressing the *acpP* guide for CRISPRi, psgRNA-*acp* was generated by inverse PCR using the primers ebm1822 and ebp8 (Table 3) as previously described (24). Amplified fragments corresponding to whole plasmids (2580 bp) were then phosphorylated (T4 Polynucleotide Kinase, Biolabs) at their 5’ extremities and circularized by ligation (T4 DNA ligase, Biolabs). *Dpn*I was then used to remove the template DNA before transformation in XL1B.

For protein purification, PCR fragments (generated from OSD56-57, 1248 bp, and OSD146-147, 263 bp respectively) corresponding to IscS and ACP coding regions were cloned between *Eco*RI and *Xho*I sites of pET6his-TEV, generating pET6his-TEV-*iscS* (pSF19) and pET6his-TEV-*acpP* (pSF51), respectively. Construction of the plasmids for BACTH assays, DNA inserts encoding the proteins of interest were inserted into pT25link (pEB354) or pT18link (pEB355) using *Eco*RI/*Xho*I sites. The pT18-IscS-ISD11 plasmid was constructed using Gibson assembly using PCR-amplified IscS (primers OSD49-50), ISD11 (primers OSD51-52) and *Eco*RI/*Xho*I linearized pT18link.

Mutated alleles of *acpP* were obtained by PCR (with the PfuUltra II DNA polymerase) using primers containing the mutations: OSD266-267 for ACP_D35N_, ebm114-115 for ACP_S36T_, OSD268-269 for ACP_D38N_ and OSD270-271 for ACP_E41Q_ (see Table 3 for sequences), and the plasmids with the wild type *acpP* allele as template. After amplification, template DNA was removed using the *Dpn*I restriction enzyme.

### Co-purification assay

Co-purifications on calmodulin beads were mainly performed as previously described (51). We worked with soluble extracts of MG1655 strain carrying pBAD-CBP derivatives (CBP-ACP fusions with either the wild type allele or the mutated alleles of ACP). Extracts were prepared from 100 mL of cultures induced by arabinose 0.01% for 1 h. Bacterial pellets were sonicated in 4 mL of calmodulin binding buffer (10 mM Tris-HCl pH 8.0, 150 mM NaCl, 0,1% NP40, 1 mM Mg-acetate, 1 mM imidazole, 2 mM CaCl_2_). After centrifugation (30 min, 20,000 g, 4°C), supernatants (3 mL) were applied on calmodulin beads (40 µL) and incubated for 2 h at 4°C. the beads were then washed three times in 1 mL of calmodulin binding buffer, then resuspended in 50 µL of Laemmli loading buffer, and then heated 5 min at 95°C. After centrifugation to eliminate the beads, extracts were loaded on SDS-PAGE for Coomassie blue coloration and Western-blot detection of IscS using polyclonal antibodies.

### mRNA extraction

Strains were grown in LB-Amp-Cm supplemented with 0.1 ng/mL AnTet at 37°C with aeration until OD_600nm_ = 1. Cells were pelleted by centrifugation (10 mL) and immediately frozen at −20 °C. RNA extractions were then performed as previously described (52). Briefly, bacterial pellets were resuspended in a buffer containing glucose 20%, Tris-HCl 25 mM, pH 7.6 and EDTA 50 mM. Cells were broken in presence of glass beads (0.1 mm diameter) and acid phenol (pH 4.5) in a cell disruptor. Successive purification steps with Trizol and chloroform were performed before isopropanol precipitation. Purified RNAs were then treated with DNase I by using the TURBO DNA-free reagent (Ambion) in order to eliminate residual contaminating genomic DNA.

### Gene expression analysis by quantitative qRT-PCR

cDNA synthesis was carried out as previously described (53). Oligonucleotides were designed in order to synthesize 100-200 bp amplicons (see Table 3). Quantitative real-time PCRs (qRT-PCRs), critical threshold cycles (CT) and *n*-fold changes in transcript levels were performed and determined using the SsoFast^TM^ EvaGreen Supermix (Bio-Rad) and normalized with respect to 16S rRNA whose levels did not vary under our experimental conditions. We analyzed the results using the Bio-Rad CFX Maestro software. Assays were performed using quadruplicate technical replicates and repeated with three independent biological samples. Results are presented as the means of the technical replicates and error bars are the standard errors of the means. Biological replicates were treated independently and did not show any significant variations.

### HPLC analysis of tRNA modifications

To extract total tRNAs, *E. coli* cells (1 L) were grown until OD_600nm_ = 1.5 and tRNA were extracted as previously described (54). Total purified tRNA (100 mg) was digested by nuclease P1 with a ratio of 2 units nuclease / 100 µg of tRNA, overnight at 37°C, followed by alkaline phosphatase treatment. Hydrolyzed tRNAs were analyzed by HPLC as previously described (55). Briefly, digested tRNAs were injected onto a C18 reverse phase HPLC column (Kinetex 4.6 x 250, 100 Å, 5 µm, packing, Phenomenex) pre-equilibrated with 100% Solvent A (2.5 % methanol and 10 mM NH_4_H_2_PO_4_, pH5.3). Nucleosides were eluted using Solvent B (20% methanol and 10 mM NH_4_H_2_PO_4_, pH5.1) and Solvent C (35% acetonitrile and 10 mM NH_4_H_2_PO_4_, pH4.6) using a gradient. Nucleosides were detected by following the absorbance at 260 nm and the retention time of each desired modified nucleosides such as s^2^C, mnm^5^s^2^U, s^4^U and ms^2^i^6^A was determined using a characteristic UV spectrum as previously described (55). The levels of each modified nucleoside are estimated based on the area of each modified nucleoside relative to 100 mg of total tRNAs.

### ß-galactosidase assay

Cells were grown in LB medium at 37°C in biological triplicates. ß-gal activity was determined as previously described (49). Average values of ß-gal unit/mg of bacteria are represented and error bars correspond to the standard deviation of the means.

### Bacterial Adenylate Cyclase Two-Hybrid assay (BACTH assay)

We used BACTH to test protein-protein interactions as previously described (20, 56). After co-electroporation of the BTH101 strain with the two plasmids expressing the hybrid proteins, cells were spread on LB-Amp-Kan plates. Liquid cultures of clones grown in LB-Amp-Kan-IPTG (0.5 mM) were then spotted on LB-Amp-Kan plates with IPTG 0.5 mM and X-Gal 50 µg/mL plates and incubated at 30 °C for two days and scanned (SCAN 4000, Intersciences).

### Gentamycin killing assay

Strains were grown aerobically in LB at 37 °C to an OD_600nm_ of 2. Cultures were then diluted in LB for an equivalent OD_600nm_ = 1. At this point, gentamycin (5 µg/mL) was added to the cells. Aliquots were taken from the culture at indicated time points, diluted in phosphate buffered saline solution (PBS), and colony-forming units (CFUs) were determined. The CFUs at time-point 0 (used as the 100 %) was around 10^9^ CFU/mL in all experiments.

### 6His-Tagged proteins purification

Recombinant 6His-IscS and 6His-ACP proteins and their derivatives were purified. Briefly, MG1655(DE3)/pET-6His-TEV-*acpP* and MG1655(DE3)/pET-6His-TEV-*iscS* strains were grown in LB (1 L) at 37°C until OD_600nm_ = 0.5, and then, 1 mM IPTG was added to induce fusion protein gene expression. For MG1655(DE3)/pET-6His-TEV-*acpP* and derivatives cultures, 10 mM of pantothenate was added to the culture (57). Cultures were further incubated at 30°C for 4 hours. After pelleting, cells were broken by sonication in 40 mL of buffer A (100 mM Hepes, 150 mM NaCl) and the extract were then centrifuged at 30,000 g for 45 min at 4°C. Supernatants were loaded onto a 5 mL Ni-nitrilotriacetic acid agarose column equilibrated with buffer A. The column was washed with 20 volumes of buffer A, and the proteins were eluted with an imidazole gradient (30 to 500 mM). Fractions were pooled and dialyzed overnight at 4°C against buffer A to remove imidazole. Purified recombinant protein quality was checked by SDS-PAGE analysis and quantified using a nanodrop and cognate epsilon values.

### *In vitro* acylation of ACP

The tagged acyl-ACP synthetase, 6His-Aas, was purified as previously described using the pET28-Aas-6his plasmid (pEB0817) (58). Acylation reactions were carried out by incubating 1 µg of purified ACP with 20 µg of palmitate and 0.25 µg of Aas in an acylation buZer (100 mM Tris-HCl pH8, 0.5 mM DTT, 10 mM ATP, 10 mM MgSO_4_) in a total volume of 20 µL. The reactions were incubated at 37°C during 3 h. The resulting ACP species were then analyzed on a 15% SDS-PAGE stained with Coomassie Blue (23).

### *In vitro* IscS-ACP binding assay

Purified 6His-IscS (15 µM) was incubated with purified 6His-ACP (30 µM) or the ACP variant 6His-ACP_D35N_ and 6His-ACP_S36T_ (30 µM) in a 10 µL reaction mix in purification buffer A (100 mM Hepes, 150 mM NaCl). Reac4ons were incubated at 37°C for 20 min before migra4on on a non-denaturing 12 % PAGE and staining with Coomassie Blue.

### Cysteine desulfurase enzymatic activity assay

IscS cysteine desulfurase activity was determined using the methylene blue assay as previously described (59). Briefly, sulfide produced by IscS desulfurase activity was monitored as follows. Cysteine (1 mM) was added to purified proteins and the mix was incubated for indicated times at 37°C in presence of dithiothreitol (DTT) 2 mM. Reactions were quenched by the addition of *N,N*-dimethyl-*p*-phenylenediamine (DMPD) 2.5 mM and FeCl_3_ 3 mM. After 30 minutes of incubation at 37°C, reactions were centrifuged 5 min at 20,000 g in order to eliminate protein aggregates. Methylene blue formation was monitored at 670 nm in a spectrophotometer. Sodium sulfide, Na2S, was used as a standard for calibration.

### Aconitase enzymatic activity assay

Aconitase activity was measured by following transformation of isocitrate to cis-aconitate, which can be monitored at 240 nm in a UV spectrophotometer as previously described (60). Briefly, cells were grown in LB with Amp, Kan and AnTet (0.1 ng/mL) when specified until OD_600nm_ around 2. Spheroplasts were immediately prepared by incubating cells (around 2.10^9^) in Tris buffer 25 mM pH 7.8 with sucrose 0.5 M on ice for 10 min and a further lysozyme treatment (0,2 mg/mL). Spheroplasts were kept at −20°C and treated individually to avoid any time laps before aconitase test. After sonication, extracts were immediately incubated in presence of isocitrate as substrate (50 mM Tris pH7.8, 0.5 mM MnCl_2_, 20 mM isocitrate) at 30°C, and absorbance at 240 nm was followed during 2 min. For each extract, protein concentration was determined by a Bradford assay. Specific activities were calculated using an extinction coefficient of 3.6 mM^-1^ cm^-1^ for cis-aconitate.

### Modeling the ACP_2_-IscS_2_ complex structure with 4’-PP and PLP modifications

The structure of the ACP_2_-IscS_2_ complex was modeled using Boltz-1 (19). Boltz-1 uses multiple sequence alignment (MSA) of the protein sequences to model as input and we used the automatic MSA generation implemented in Colabfold (61). We modeled the complex with the 4’-PP bound to the serine 36 of the ACP chains, and the PLP bound to the lysine 206 of the IscS chains. For the 4’-PP, we used the “modification” field of the yaml configuration file of Boltz-1 with the CCD code 4HH (https://www.rcsb.org/ligand/4hh) corresponding to the 4’-PP-serine. Since no CCD is available on the Protein Data Bank (PDB) for the PLP bound to lysine, we add a new residue coding the PLP bound to lysine into the CCD cache of Boltz-1 as described in https://github.com/benf549/boltz-generalized-covalent-modification/tree/main. We named KPLP the new CCD encoding the PLP bound to lysine. The scripts used to add the KPLP CCD to the CCD cache of Boltz-1 is given in supplementary materials, as well as the yaml configuration file of Boltz-1.

## Acknowledgements

We are grateful to Béatrice Py, Sandrine Ollagnier de Choudens and Benoît d’Autréaux for helpful discussions. We also thank all the members from the SAMe unit for feedback and suggestions. This work was supported by the ANR-10-LABX-62-IBEID, the Institut Pasteur and the CNRS. S.F. received a PhD fellowship from Université Paris-Cité.

## Supplementary informations

### Supporting information text 1 Boltz-1 scripts

**Scripts used to add the KPLP CCD to the CCD cache of boltz-1:**

– modified_residues.py: Python scripts to add a new residue into the CCD cache of boltz-1: https://github.com/bougui505/misc/blob/master/singularity/boltz/modified_residues.py. The command line to run the script is:1

./modified_residues.py-s “CC1=NC=C(COP(O)(O)=O)C(CNCCCCC(N)C(O)=O)=C1O” Ä -a LYS -o KPLP

**ACP2holo_PLP_IscS2.yml: yaml configuration file of boltz-1 to build the ACP_2_-IscS_2_ complex structure with 4’PP and PLP** modifications:

sequences:

– protein:

id: A

sequence: STIEERVKKIIGEQLGVKQEEVTNNASFVEDL GADSLDTVELVMALEEEFDTEIPDEEAEKITTVQAAIDYINGHQA

modifications:

– position: 36 ccd: ‘4HH’

– protein:

id: B

sequence: STIEERVKKIIGEQLGVKQEEVTNNASFVEDL GADSLDTVELVMALEEEFDTEIPDEEAEKITTVQAAIDYINGHQA

modifications:

– position: 36 ccd: ‘4HH’

– protein:

id: [C, D]

sequence: MKLPIYLDYSATTPVDPRVAEKMMQFMTMDGTFGNPASRSHRF GWQAEEAVDIARNQIADLVGADPREIVFTSGATESDNLAIKGAANFYQKKG KHIITSKTEHKAVLDTCRQLEREGFEVTYLAPQRNGIIDLKELEAAMRDDT ILVSIMHVNNEIGVVQDIAAIGEMCRARGIIYHVDATQSVGKLPIDLSQLK VDLMSFSGHKIYGPKGIGALYVRRKPRVRIEAQMHGGGHERGMRSGTLPVH QIVGMGEAYRIAKEEMATEMERLRGLRNRLWNGIKDIEEVYLNGDLEHGAP NILNVSFNYVEGESLIMALKDLAVSSGSACTSASLEPSYVLRALGLNDELA HSSIRFSLGRFTTEEEIDYTIELVRKSIGRLRDLSPLWEMYKQGVDLNSIE WAHH

modifications:

– position: 206 ccd: ‘KPLP’

---

**Fig. S1:**
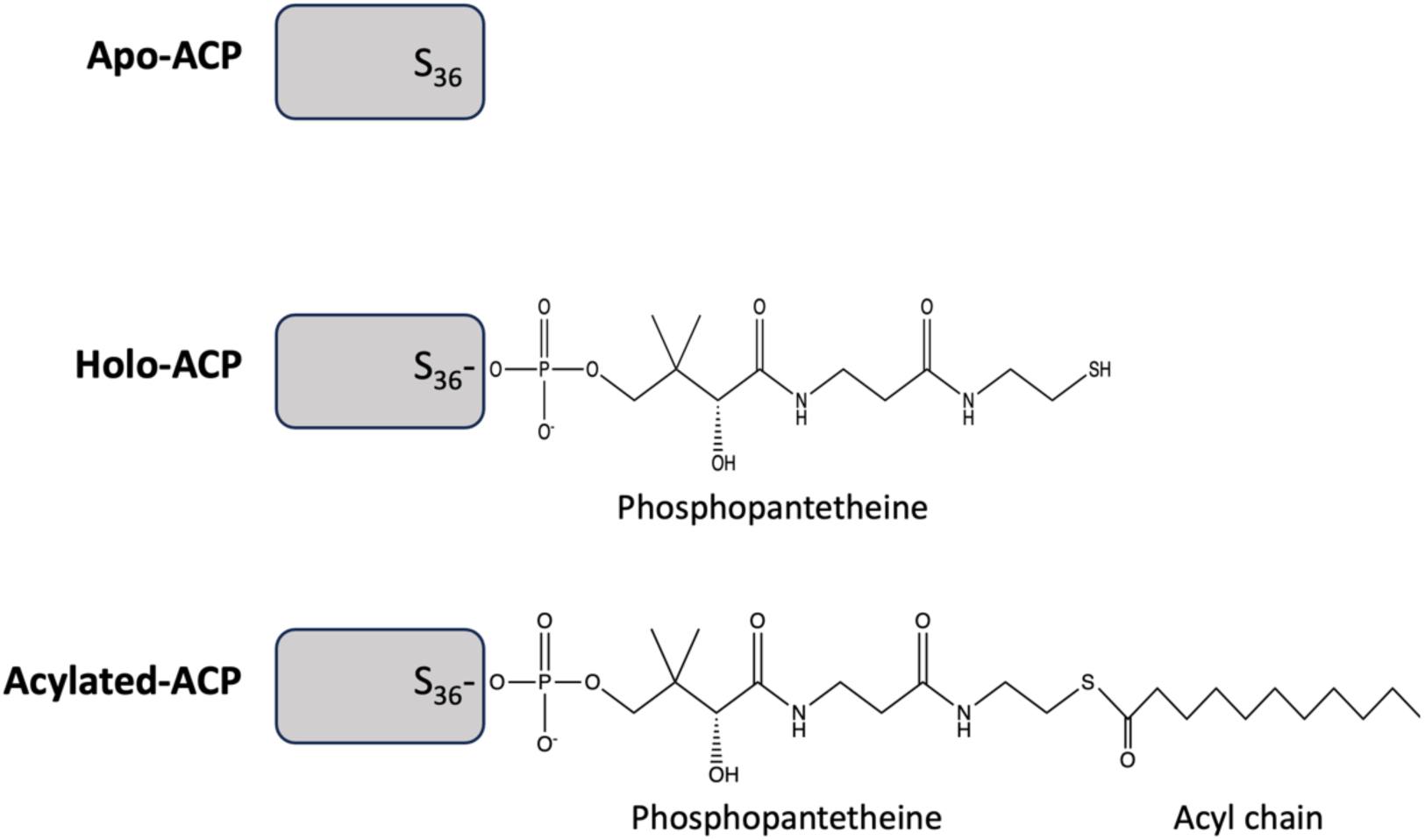
The three molecular species of ACP found in the cell. ACP is initially synthesized in its apo­ form. The first maturation step involves the covalent attachment of a 4’-phosphopantetheine group to the hydroxyl group of serine 36 in ACP via a phosphodiester bond, converting it to holo­ ACP. In the second step, a malonyl group is transferred to the terminal thiol group of phosphopantetheine through transesterification. This loaded ACP then transports the malonyl chain to enzymes in the FASII pathway, where it undergoes elongation. The nature of the acyl chain linked to ACP varies depending on the stage of elongation.

**Fig. S2:**
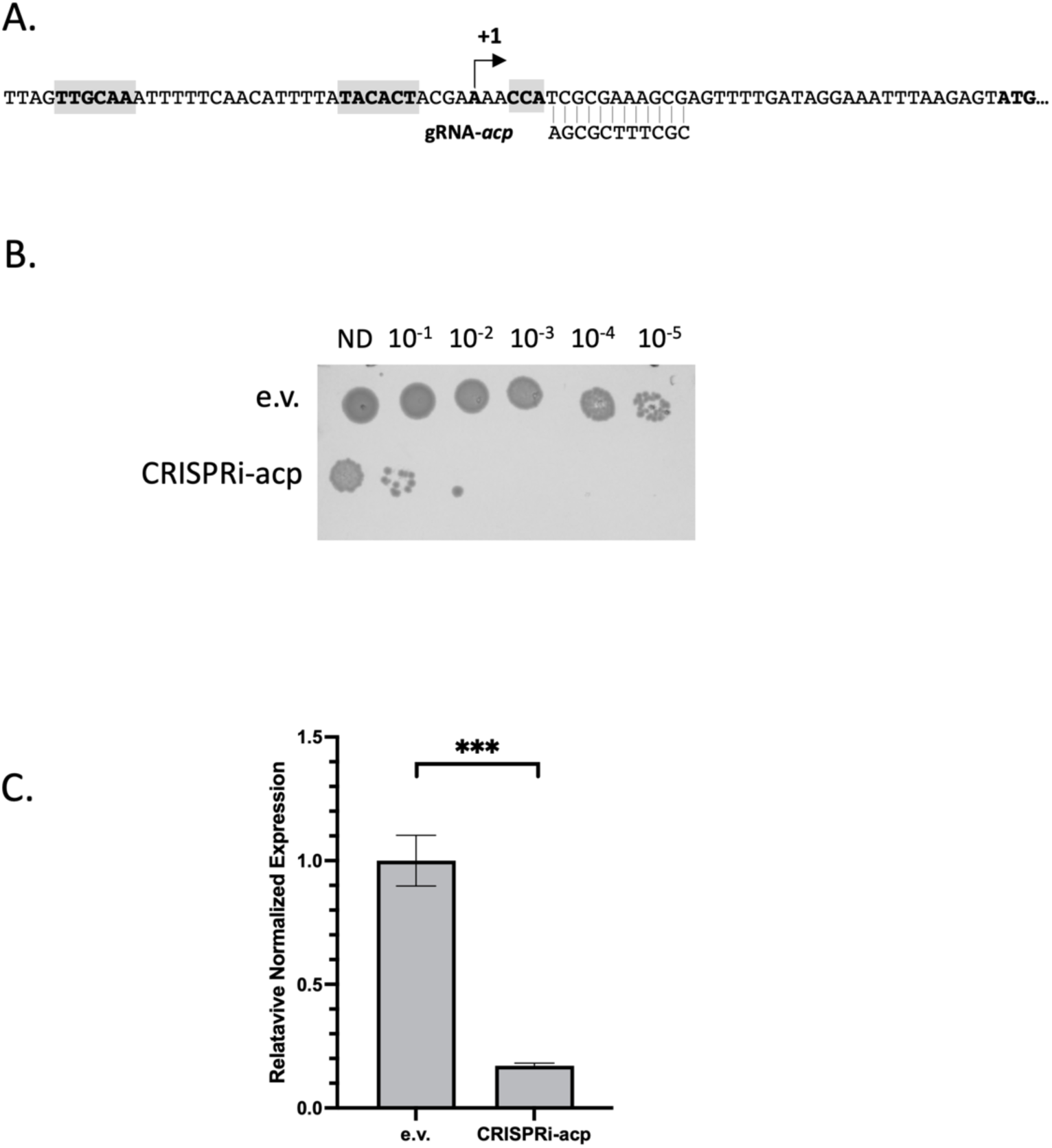
Validation of the CRISPRi tool to decrease *acpP* expression. **A.** Upstream region of *acpP* coding sequence. Grey boxed sequences upstream the transcriptional start site (+1) indicate −10 and −35 boxes. Grey boxed sequence downstream the +l indicates the PAM sequence located in front of the complementary sequence of the guide RNA we used (gRNA-acp). **B.** MG1655 WT strain was co­ transformed with either the pdCas9 and the psgRNA plasmids (e.v.) or with the pdCas9 and psgRNA­ *acp* plasmids (CRISPRi-acp). Cells were grown in LB until OD_6_aanm = 1 and indicated dilution were spotted (3 µL) on LB plate containing 500 ng/mL AnTet in order to induce dCas9 expression. Plates were incubated one night at 37°C before imaging. **C.** Same strains as described above were cultivated in LB at 37°C until OD_60_anm = 1 in presence of 0.1 ng/mL of AnTet. RNA were extracted and treated as described in the Materials and Methods section. Quantitative Real-Time PCR was used to quantify *acpP* cDNA derived from RNA. Expression levels were normalized using 165 rRNA as internal standard and presented as the n-fold change of the CRISPRi-acp strain compared to the e.v. strain. Results are presented as the mean of4 replicates and standard errors to the means are indicated.***, P<0.005.

**Fig. S3:**
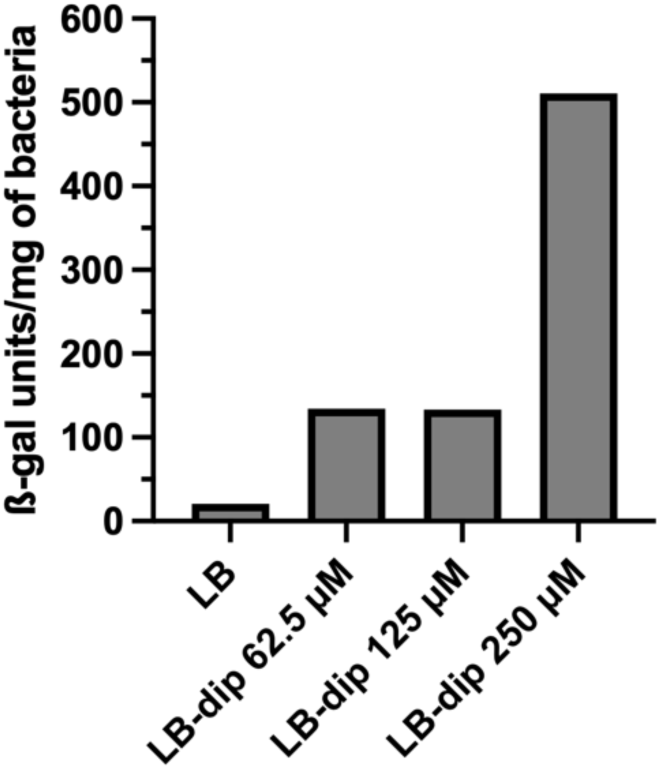
Study of the *iscR* expression under iron depletion. Bacterial cells, *PiscR-/acZ* strain carrying the CRISPRi plasmids, pdCas9 and psgRNA (e.v.), were inoculated in LB with increasing concentrations of 2-2’-dipyridyl (DIP) until OD_6_aonm = 2. Activity of the *iscR* promoter was assessed by measuring the B-gal activity (Miller units). We noticed that the presence of DIP 62.5 and 125 µM did not alter bacterial growth.

**Fig. S4:**
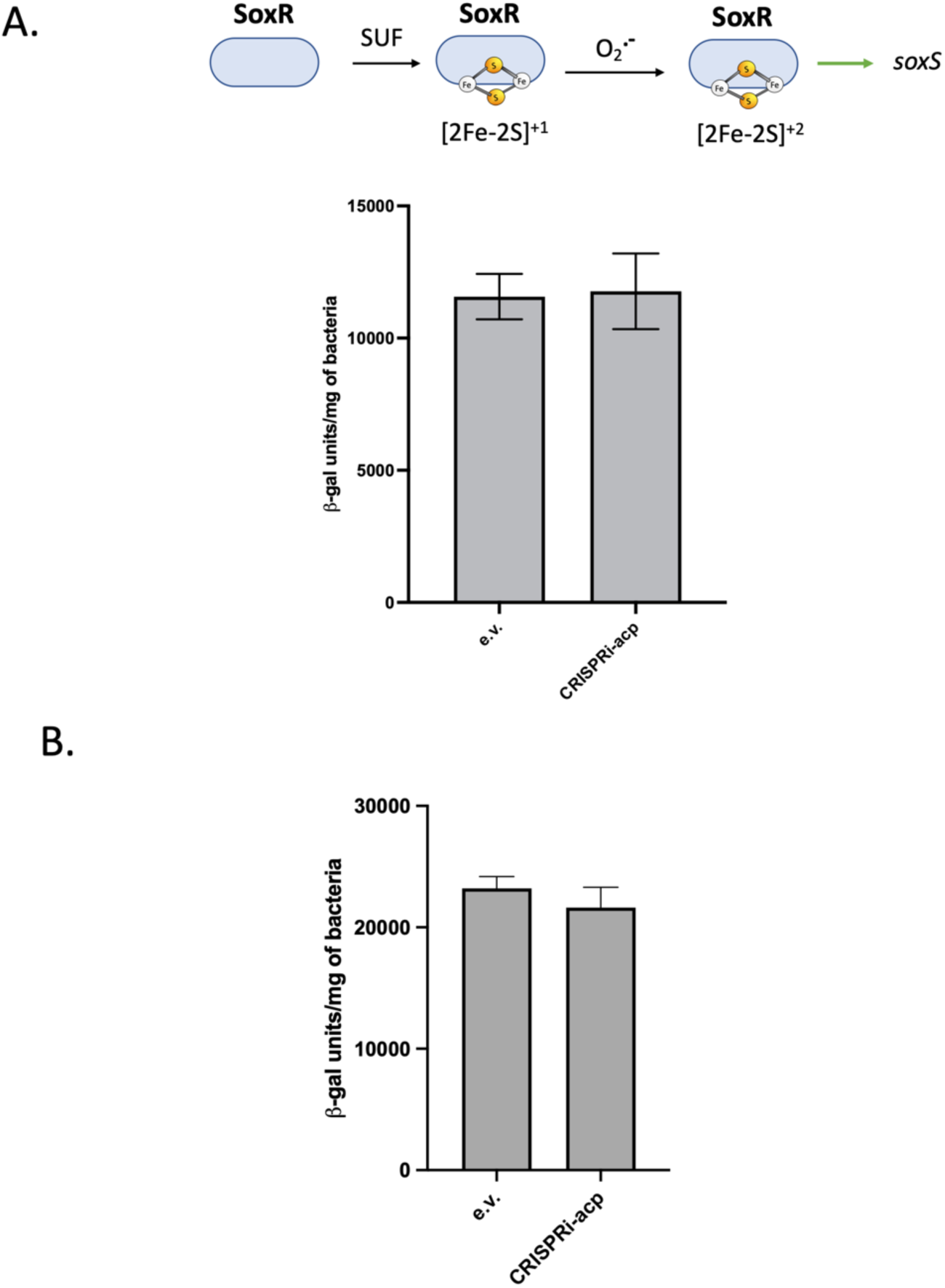
ACP does not affect the activity of proteins not matured by the ISC system. A. Effect of ACP levels on SoxR-dependent regulation under oxidative stress condition. Activity of the *SoxR-activated soxS* promoter was assessed by measuring the f3-gal activity (Miller units) of the *Psox-lacl* strain carrying the indicated plasmids in a WT (MG1655) background. Strains were grown in LB with O.lng/mL of AnTet until _00600nm_ = 1 and then treated with paraquat dichloride (100 µM) for 1 hour before B-gal activity was measured. **B. Effect of ACP levels on endogenous f3-galactosidase activity.** Wild type MG1655 strain carrying indicated plasmids have been grown in LB with 0.lng/mL of AnTet and 0.5mM of IPTG until _00600nm= 2._ Represented values are the mean of three independent biological replicates and the error bars represent the standard error to the mean.

**Fig. S5:**
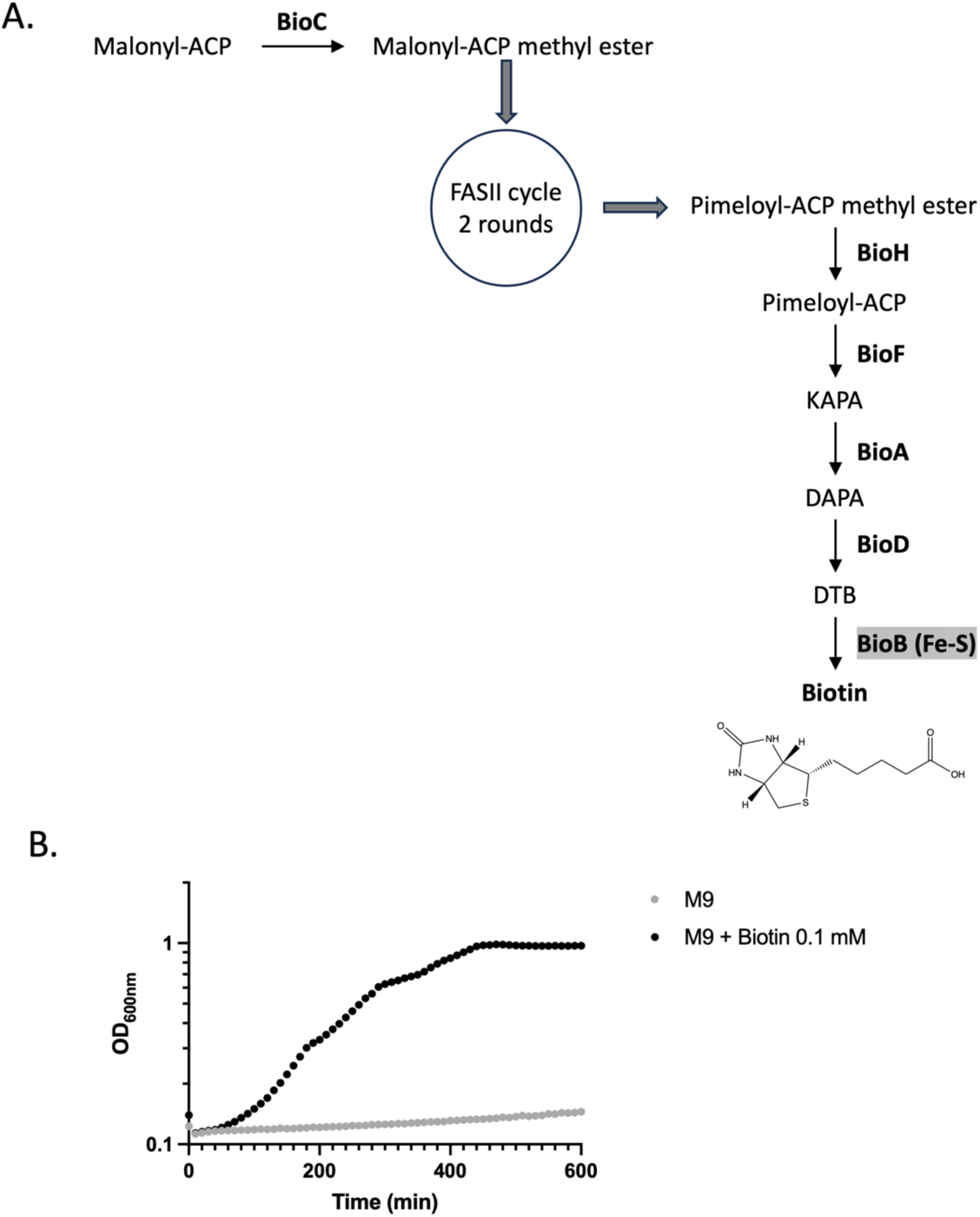
Biotin is synthesized through a complex pathway from a fatty-acid bound ACP precursor and is essential for bacterial viability. A) *E. coli* biotin biosynthetic pathway. BioB is the radical SAM Fe-5 enzyme that catalyses the last step of biotin synthesis by incorporation of a sulfur atom in dethiobiotin (DTB). KAPA: 7-keto-8-amino-pelargonic acid; DAPA: 7,8-diaminopelargonic acid. **B)** Growth of MG1655 *flbioD* strain in M9 minimal medium with glucose 0.2 % as carbon source and supplemented (black points) or not (grey points) with 0.1 mM biotin.

**Fig. S6:**
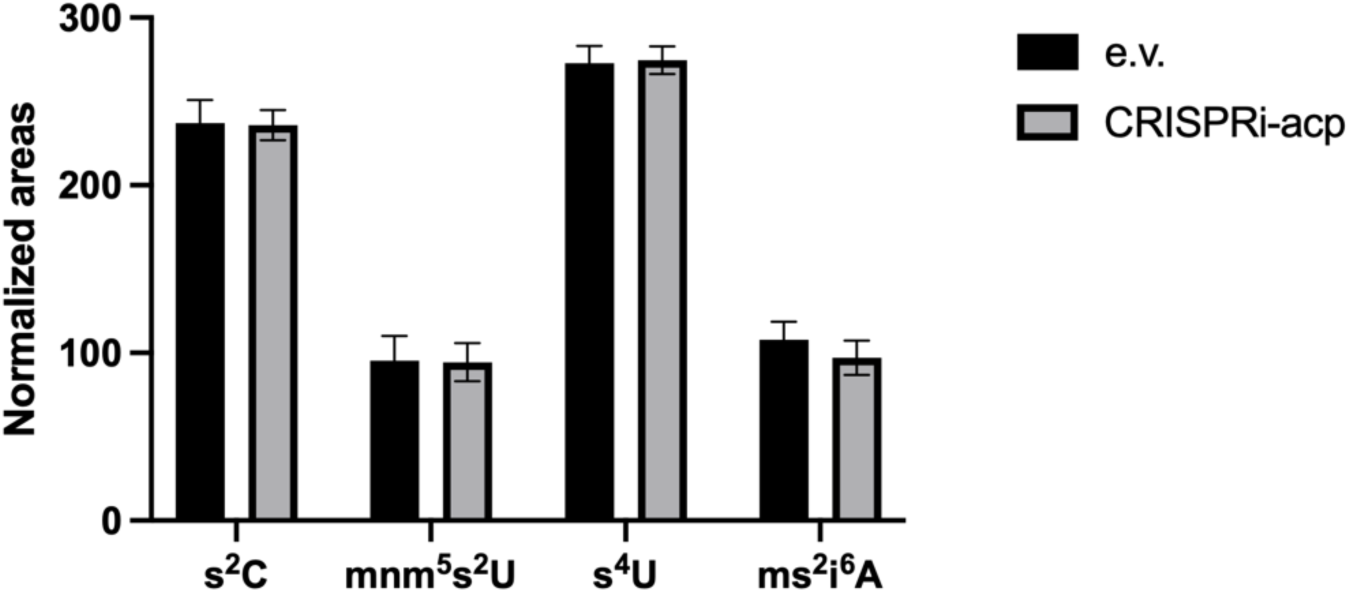
ACP levels do not impact lscS-dependent tRNA thiolation. Relative levels of specific tRNA modified nucleosides in bacterial cells (MG1655) carrying pdCas9 and psgRNA (e.v., black bars) or psgRNA-ocp (CRISPRi-acp, grey bars) grown in LB with 0.1ng/ml of AnTet. The retention time of modified nucleosides of s^2^C, mnm^5^s^2^U, s^4^U and ms^2^i^6^A was determined using a characteristic UV spectrum as previously described (1). The levels of modified nucleosides are estimated based on the area of each modified nucleoside relative to 100 mg of total tRNAs. The data are presented as the mean of four replicates and and the error bars represent the standard error to the mean. 1. C. W. Gehrke, K. C. Kuo, Ribonucleoside analysis by reversed-phase high-performance liquid chromatography. *J Chromatogr471,* 3-36 (1989).

**Fig. S7:**
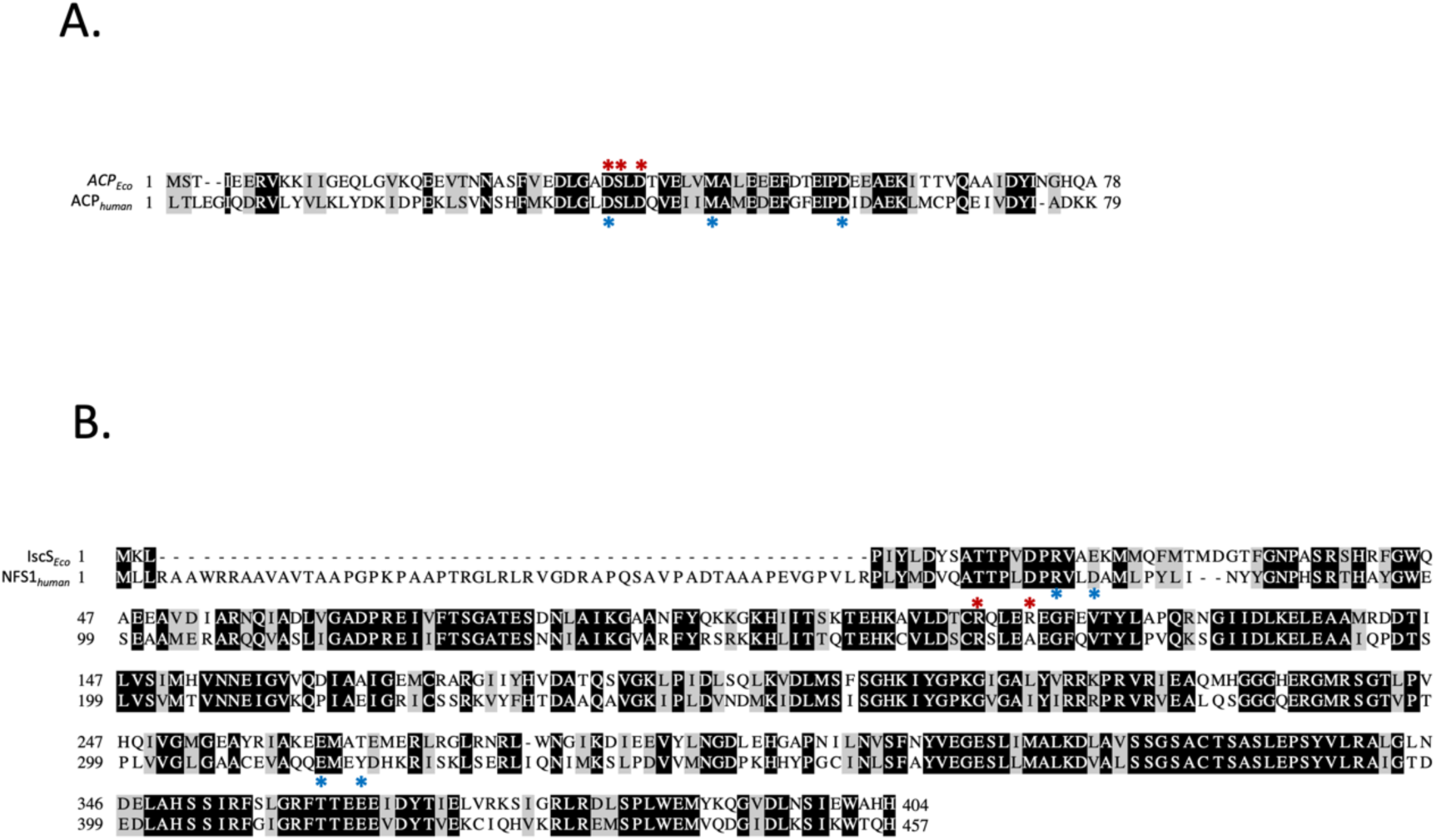
Comparison of primary sequences of ACP and lscS from *E. coli* and human. **A.** Amino acid sequence alignment of ACP from f. *coli* (first line) and human ACP (second line) and **B.** lscS from *E. coli* (first line sequence) and Human Nfsl (second line). Black boxes indicate identical residues, and grey ones, similar residues. Red stars indicate amino acids (for ACP: D_35_, S_36_, D_38_ and for lscS: R_112_, and R_116_) thought to be involved in ACP-lscS interaction (this work). Blue stars indicate human ACP and NFS1 residues involved in interaction with 15D11 (D_35_, M_44_, D_56_ and R_72_, D_75_, E_31_,., Y_317_ respectively) (1). 1. M. T. Boniecki, S. A. Freibert, U. Muhlenhoff, R. Lill, M. Cygler, Structure and functional dynamics of the mitochondrial Fe/S cluster synthesis complex. *Nat Commun* **8,** 1287 (2017).

